# From Bulk to Binding: Decoding the Entry of PET into Hydrolase Binding Pockets

**DOI:** 10.1101/2024.04.21.590457

**Authors:** Anna Jäckering, Frederike Göttsch, Moritz Schäffler, Mark Doerr, Uwe T. Bornscheuer, Ren Wei, Birgit Strodel

## Abstract

Plastic-degrading enzymes hold promise for biocatalytic recycling of poly(ethylene terephthalate) (PET), a key synthetic polymer. Despite their potential, the current activity of PET hydrolases is not sufficient for industrial use. To unlock their full potential, a deep mechanistic understanding followed by protein engineering is required. Using cutting-edge molecular dynamics simulations and free energy analysis methods, we uncover the entire pathway from the initial binding of two PET hydrolases – the thermophilic leaf-branch compost cutinase (LCC) and polyester hydrolase 1 (PES-H1) – to an amorphous PET material to a PET chain entering the active site and adopting a hydrolyzable geometry. Our results reveal the initial PET binding and elucidate its non-specific nature driven by electrostatic and hydrophobic forces. Upon PET entry into the active site, we uncover that this process can occur via one of three key pathways and detect barriers to it arising from both PET–PET and PET–enzyme interactions, with specific residues identified by *in silico* and *in vitro* mutagenesis. These insights not only advance our understanding of PET degradation mechanisms and pave the way for targeted enzyme enhancement strategies, but also offer an innovative approach applicable to enzyme studies across disciplines.

## 1 Introduction

The plastic consumption of our society exhibits an escalating trajectory with each passing year, accompanied by a commensurate escalation in the volume of plastic waste that necessitates recycling.^1^ However, the efficiency of plastic recycling remains suboptimal, in particular for composite materials. But even when dealing with a high degree of homogeneity in a given plastic, the employed recycling methodologies are invariably accompanied by significant greenhouse gas emissions. ^2^ For this reason, the quest for an environmentally-friendly enzymatic depolymerization process of plastic waste that allow for the monomer recovery has attracted attention, especially for the widely used polymer poly(ethylene terephthalate) (PET).^3–6^

PET is an aromatic, semicrystalline thermoplastic that can be enzymatically degraded into monomeric and oligomeric building blocks including 2-hydroxyethylterephthalic acid (MHET), which can then be further hydrolyzed into its monomers terephthalic acid (TPA) and ethylene glycol (EG) by the same or different enzymes. ^4,7–9^ The monomers can be recovered to synthesize high quality virgin PET (Fig. 1).^3,10^ In 2005, enzymatic degradation of low-crystallinity PET, which resulted in a significant mass loss, has been reported for the first time with a *Thermobifida fusca* hydrolase.^11^ Afterwards, many other ester hydrolases (EC 3.1.1.x) were found to be capable of degrading PET, which were assigned to the novel subclass for PETases (EC 3.1.1.101).^12^ A milestone in microbial PET degradation was achieved eleven years later with the discovery of *Ideonella sakaiensis*, which features a two-enzyme system for the depolymerization of PET and its degradation products as the main source of carbon and energy for the bacterial growth. ^13^ However, the activity and thermostability of wild-type (WT) PET hydrolases are often insufficient for their industrial application. To this end, diverse PET hydrolases such as the leaf-branch-compost cutinase (LCC) and the polyester hydrolase 1 (PES-H1) have been recently engineering yielding two of the currently most active and thermostable PET hydrolases, the LCC F243I/D238C/S283C/Y127G (ICCG) variant^3^ and the PES-H1 L92F/Q94Y (FY) variant.^14^ In particular, the development of LCC^ICCG^ by the French company Carbios enabled the launch of a first enzymatic PET recycling demonstration plant, initiating the evaluation of its pre-industrial-scale recycling process, which involves 100 tons of pretreated PET waste per year.^3,15,16^ The increased thermal stability of LCC^ICCG^ allowed for its application in the temperature range of 68-72°C, where the PET polymers gain high mobility and are easily accessible to the enzymes.^3,4,17–19^ At the same time, amorphous PET can recrystallize quickly in the aqueous milieu at temperatures above 72°C, necessitating rapid kinetics of enzymatic PET degradation when used in this temperature range, which remains a significant challenge for enzyme engineering.^3^

**Figure 1:**
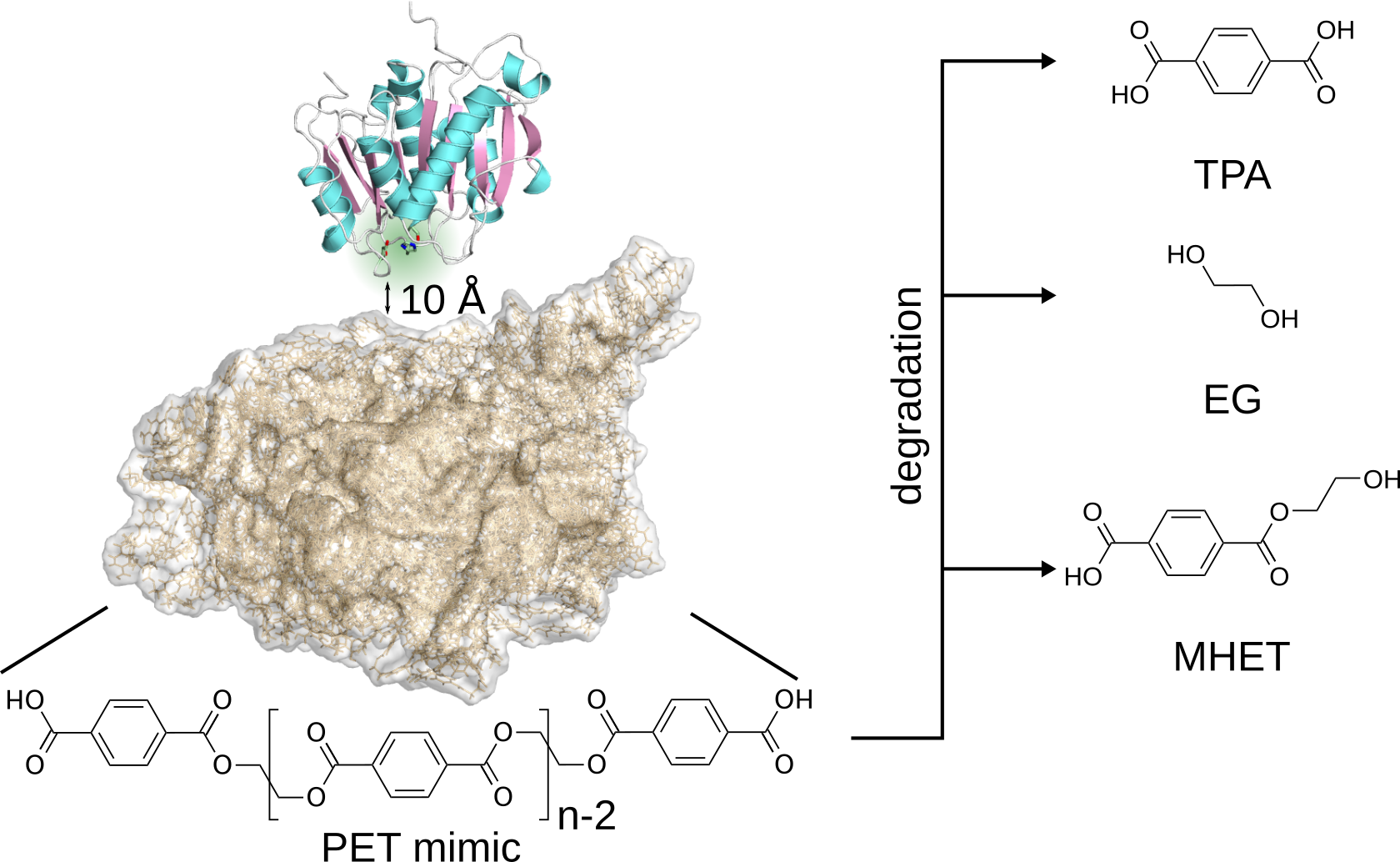
Representative starting structure for the MD simulations of enzyme adsorption on amorphous PET (without water and ions being shown). The system comprises one PET hydrolase, which is shown as cartoon and with the catalytic triad highlighted in green. The enzyme was randomly oriented and placed at least 10 Å away from the PET bulk, which is shown as brown surface and consists of 100 PET-9mers. The chemical structure of PET is shown at the bottom and the degradation products of PET, terephthalic acid (TPA), ethylene glycol (EG), and 2-hydroxyethyl terephthalic acid (MHET), are shown on the right.

To make further progress here, a deeper understanding of the entire PET degradation process is required. This process consists of initial adsorption of the enzyme to the PET bulk, followed by a single PET chain from this bulk entering the active site of the PET hydrolase and assuming a binding position productive for subsequent hydrolysis. It was suggested that the PET hydrolases adsorb non-specifically on the surface of the PET bulk, resulting in either a productive complex that enables PET degradation by binding close to the active site, or in an unproductive complex that cannot hydrolyze PET.^20,21^ Therefore, previous engineering studies have focused on modifying general surface properties such as charge, polarity and hydrophobicity of the PET hydrolases to improve PET adsorption. ^22^ Increased PET adsorption was observed after the introduction of positive charges ^23,24^ and increased hydrophobicity of the enzyme surface. ^25–28^ However, according to the Sabatier principle, optimal catalysis occurs when interactions between catalyst and substrate are of intermediary strength. For LCC it was demonstrated that its activity increased when its substrate affinity is weakened due to addition of a cationic surfactant.^29^ Thus, the optimal balance between binding affinity and activity of PET hydrolases remains unclear, necessitating a thorough examination of the initial PET adsorption process and the transition from non-specific adsorption to a productive PET binding mode at the active site. Therefore, it is not yet clear how to optimize the relationship between binding affinity and activity of PET hydrolases, which requires a detailed analysis of the initial PET adsorption and transition from adsorption far from the active site to productive PET binding.

Most design approaches target the formation of the substrate–enzyme complex that takes place in the catalytic binding cleft to enhance the activity of PET hydrolases. Many studies have introduced substitutions inspired by comparing the amino acid composition of different PET hydrolases in the active site, resulting in the PES-H1^FY^ variant, which was motivated by the thermostable DuraPETase.^14,30^ Computational methods offer insights into the importance of specific residues in the binding site, which complements the limited high-resolution crystal structures of PET hydrolase with small ligands reported recently.^9,14,23,31–35^ Based on the analysis of these structural complexes, it was hypothesized that widening the binding cleft would increase the accessibility of the active site and consequently the enzyme’s activity,^36–39^ while narrowing the binding site could enable specific interactions with PET that promote substrate binding and enzyme activity.^40^ Since PET is an aromatic, mostly hydrophobic polymer that can form hydrogen bonds, it has been shown that the introduction of aromatic, hydrophobic or hydrogen bond-forming residues also has a positive effect on the efficiency of PET hydrolases. ^37,39,41,42^ A highly conserved tryptophan that flanks the binding cleft and appears to be crucial for enabling PET degradation deserves special mention. Its flexibility diverges between mesophilic and thermophilic enzymes that harbor different amino acids in the neighborhood of the tryptophan, which influences its ‘wobbling’ capacity.^35,43–45^ Nevertheless, not all results are consistent with respect to the factors that promote productive PET binding, indicating that further analysis is needed to identify consistent characteristics that support productive binding. Furthermore, the computational methods in the previous studies comprise mainly docking^3,35,36,46,47^ in combination with short molecular dynamics (MD) simulations. ^14,40,48–51^ However, the dynamics of the binding process, which is composed of the flexibility of the binding site itself and the PET as well as the process of PET entry into the active site, cannot be adequately described by these methods, yet they have been suggested to be important factors for PET complexation and thus degradation.^4,48,49,52–54^

To fill this knowledge gap, we performed thorough computational simulations in combination with experimental studies on two metagenome-derived thermophilic PET hydrolases, LCC (catalytic triad: S165, D210, H242) and PES-H1 (catalytic triad: S130, D176, H208). To be precise, we subjected both enzymes to extensive MD as well as Hamiltonian replica exchange MD (HREMD) simulations to elucidate their adsorption to a PET bulk, tracking PET entry into the active site until PET binds in a productive mode to allow for hydrolysis. PET entry is analyzed using a free energy surface-based approach to reveal residues that affect the energy barrier during the entry process, whereupon residue substitutions are predicted and then evaluated *in vitro*. Our approach is the first to show how PET reaches the active site, uncovering important information about the dynamics of both the enzyme and PET, as well as the interactions between polymer and enzyme.

## 2 Results and discussion

To investigate the entire interfacial biocatalysis process, starting with the enzymes LCC and PES-H1 binding to a PET bulk and ending with a PET chain in the active site ready for hydrolysis, we began our study by placing both LCC and PES-H1 in random orientations and at least 10 Å away from an amorphous PET bulk. The structure of the PET bulk was based on Cruz-Chu *et al.*^55^ including 100 PET-9mers(Fig. 1). Subsequently, we conducted a total of 10.5 µs of MD simulations for each enzyme. In these simulations, we observed enzyme adsorption on the PET surface at sites proximal and distal to the binding cleft during 75% of the simulation time, but the PET failed to enter into the active site. Therefore, we continued the simulations by applying the enhanced-sampling HREMD method (accumulating further 43.47 µs of sampling per enzyme). We used only one single PET chain closest to the binding cleft allowing PET to enter the active site and adopt productive binding poses. The results of these two sets of simulations are discussed in detail below along with the wet lab studies to consolidate our *in silico* findings.

### 2.1 PET adsorption to the surface of the enzyme

#### Both enzymes readily adsorb to PET, but without a strong preference for the binding cleft

Adsorption to the PET bulk was sampled in all six of the 1.5 µs simulations for both LCC and PES-H1. In addition, the initial enzyme–PET interactions of considerable strength with energies below *−*10 kcal*·*mol^-1^ were established early in the simulations (PES-H1: 2.0–40.2 ns; LCC: 0.8–15.0 ns) and abundant contacts (defined based on the residue–PET distance requirement *≤*3 Å) were observed at the surfaces of LCC and PES-H1, suggesting that initial binding is based on general features such as charge and polarity rather than specific recognition sites.^56–58^ These initial contacts occurred almost everywhere on the surface of both enzymes and can be divided into proximal and distal binding, depending on whether PET was located within 12 Å of the catalytic serine or not. These contact areas are colored blue and red in Fig. 2A, respectively, and the regions where distal and proximal areas overlap are indicated in purple. To follow the progression of the contacts during the simulations, we compared the contact areas during the first 50 ns and the remaining simulation time. However, only limited reorientations of the enzymes on the PET surface were observed (shown in yellow for newly explored surface and in brown for overlaps with only distal or proximal in Fig. 2A); most contacts remained unchanged for 75% or more of the simulation time. In the case of proximal binding at the end of the simulations, we extended the corresponding simulations to 2 µs to see if further PET progression towards the active site would occur. The general findings for both, simulations lasting for 1.5 µs and the elongated simulations lasting for 2 µs, are that when the initial contacts were established in proximal regions, the PET contact area remained close to the active site (PES-H1: 2 of 6; LCC: 4 of 6 simulations). In cases where the initial contacts were in distal areas, negligible progression towards the active site was observed. Only in one of the PES-H1 simulations, proximal contacts could be established starting from distal ones (highlighted in yellow and overlaps with regions establishing distal or proximal contact only in orange in Fig. 2A). The observed contact progression towards proximal binding in one of the PES-H1 simulations might contribute to the high activity of this enzyme.

**Figure 2:**
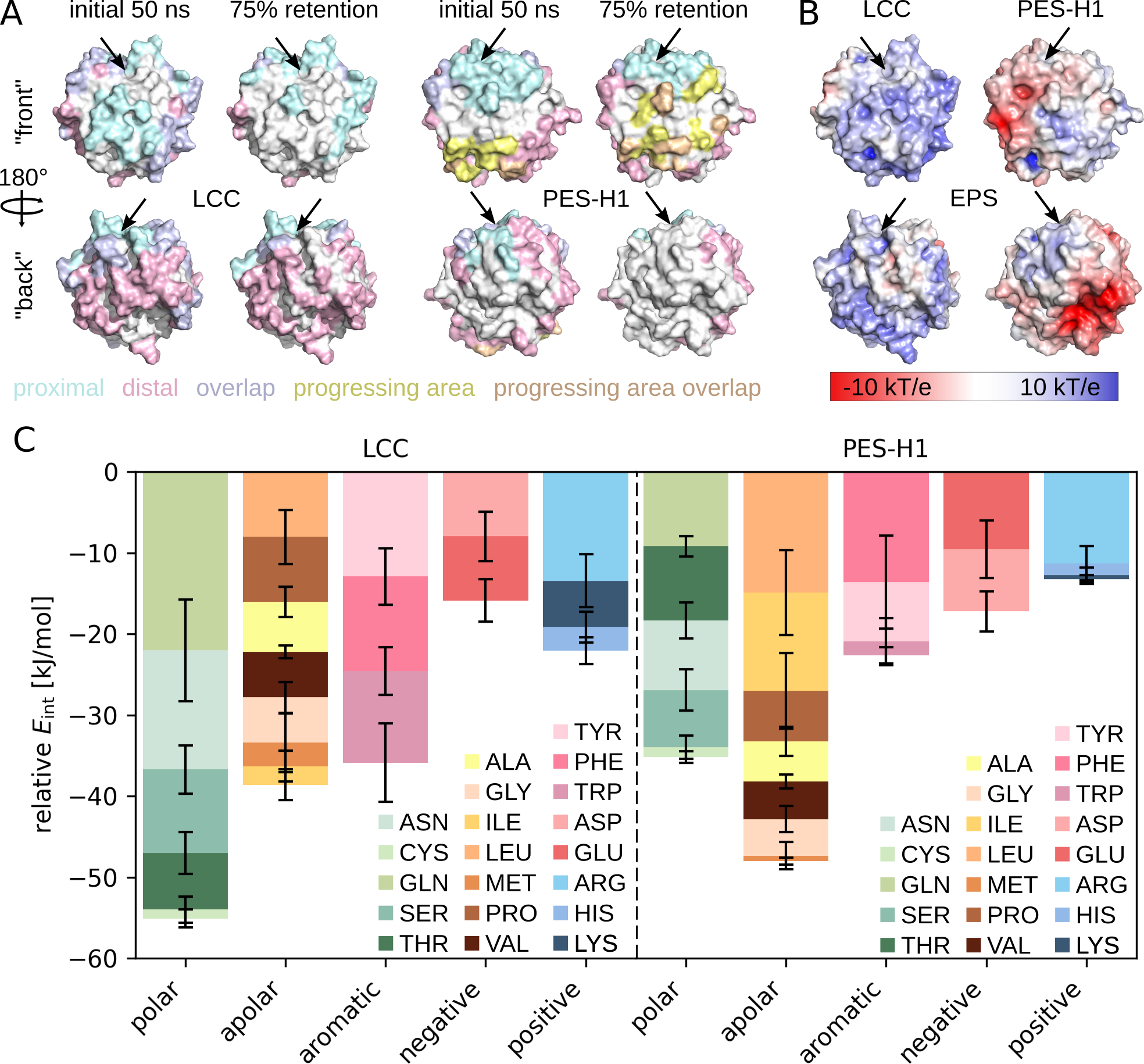
Analysis of LCC (left) and PES-H1 (right) adsorption on amorphous PET. (A) Residues contacting PET are highlighted on enzyme surfaces, using all six simulations per enzyme with a 3 Å PET–residue distance cutoff. Contacts during initial 50 ns (left) and stable contacts present for at least 75% of the full duration (right) are distinguished, with proximal (blue) and distal (rose) binding sites. Overlapping areas are colored violet. Some simulations show enzyme reorientation on PET bulk, with limited progression from distal to proximal binding, except in one PES-H1 simulation (highlighted in yellow). Overlaps with proximal or distal contact areas highlighted in brown. The position of the catalytic triad is denoted by an arrow. (B) Electrostatic potential surface (EPS) for LCC (left, net charge: +5) and PES-H1 (right, net charge: *−*5). (C) PET interaction energies *E*_int_ with amino acid types in LCC and PES-H1 are shown relative to the number of the corresponding residues within 3 Å, averaged over all simulations. Energies for amino acid types with the same physicochemical characteristics are summed up, with standard error of the mean (SEM) shown as error bars.

If this is the case, a promising way to achieve accelerated and productive PET binding could be to facilitate enhanced proximal binding while attenuating PET interactions at distal sites of the enzyme surfaces. To test this hypothesis, we first analyzed the physicochemical nature of the initial enzyme–PET contacts and then made rationally motivated mutations, which are studied *in silico* and *in vitro*.

#### Negative surface charges impede PET adsorption

LCC has a net charge of +5, while PES-H1 has a net charge of *−*5, which is reflected in their electrostatic potential surface (EPS, shown in Fig. 2B). The EPS of LCC is mostly neutral and positive (shown in white and blue, respectively), while PES-H1 features large surface patches that are strongly negative (shown in red). For PES-H1 in particular, the negative regions overlap with those that do not have pronounced PET contacts. In addition, we visualized the EPS of the PET bulk used to construct the initial enzyme–PET systems. A mostly hydrophobic surface is revealed, with predominantly convex and exposed areas being partially negatively charged, while stronger positive or negative potentials can be found in concave and buried regions (Fig. S2). We therefore assume that negative charges at the surface discourage PET binding, which could open up the possibility of preventing distal binding to PET by introducing negatively charged residues there. This idea is consistent with the results of Nakamura *et al.*, who demonstrated accelerated binding to PET surfaces with higher positive charge density at the enzyme surface of PET2.^23^ In addition, Sagong *et al.* found that changes in the electrostatic potential surface of a loop close to the active site increases PET degradation activity of *Rhizobacter gummiphilus* PETase (*Rg* PETase), especially when increasing the negative surface charge. ^59^ In order to transfer this finding from proximal regions to distal ones, they mutated four distal surface-exposed arginine to alanine and glutamate, which also increases the PET degradation activity.

This effect was more pronounced when introducing glutamate, from which they conclude that a negative surface charge is favorable for the PET degradation activity of *Rg* PETase. Moreover, Thomsen *et al.* observed that PES-H1 creates craters on the PET surface, unlike other enzymes such as LCC exhibiting a uniform degradation.^21^ They ascribe this distinct behavior in PES-H1 to its the distal negative surface patch influencing enzyme positioning and processivity after the initial contact due to repulsion with the negatively charged PET termini. We thus conclude that positive regions enhance overall PET adsorption and lead to prolonged PET retention in distal regions when present there, which in turn hinders the process of PET binding in a productive conformation at the active site.

#### Apolar residues contribute most to PET adsorption of PES-H1

To identify the amino acids most favored by PET, we calculated the interaction energy *E*_int_ between PET and each amino-acid type for all six simulations of both enzymes. We then scaled the resulting energy values by the number of residues of the corresponding amino-acid type that were in contact with PET during each simulation and took the average of the six values. These results are shown in Fig. 2C, where we grouped the amino acids according to their physicochemical property. The scaling by the contact residues accounts for the fact that more polar than apolar residues are exposed in both enzymes, making them more accessible to PET than apolar residues. The overall interaction with PET (i.e., the sum of all scaled *E*_int_ values) is stronger for LCC than for PES-H1, with polar residues contributing most to LCC binding and apolar contributing most to PES-H1 binding. Charged amino acids contribute the least to PET binding for both enzymes, followed by the aromatic ones. Given the progression from distal to proximal binding observed for PES-H1 but not for LCC in this study, we propose that modification of the surface of LCC to make it more similar to that of PES-H1 could increase proximal PET binding for LCC and thus its enzymatic activity. Based on our research findings, it appears that the PET binding is excessively strong, which leads to extended retention in unproductive adsorption orientations. Consequently, our mutation strategy aims to diminish the interactions between PET and LCC. This adjustment should facilitate the enzyme’s reorientation following its initial attachment to the PET surface. The goal is to promote closer and more favorable binding without excessively compromising the overall adsorption capability. This should be achieved by eliminating strong PET interactions with polar LCC residues and by promoting weaker interactions with apolar residues. This design strategy gains support from previous studies that revealed that enhanced activity of PET hydrolases is associated with increased enzyme hydrophobicity.^25–27^ In addition, a reduction in the hydrophobicity of LCC in proximal areas has been demonstrated to reduce the binding to a PET surface, resulting in increased enzyme activity due to increased conversion of degradation intermediates and consequently elevated TPA production. ^60^

#### Distal introduction of alanine and negative charge reduces local interaction with PET

To verify this assumption, we generated three *in silico* variants per enzyme, each containing three mutations. The mutations were incorporated into the highly active variants LCC F243I/Y127G (IG) (from LCC^ICCG 3^) and PES-H1 L92F/Q94Y (FY),^14^ which replace the WT enzymes investigated up to this point. In LCC^ICCG^ there are two other mutations which we do not consider in this part of the study. The reason for this is that removing the negative charge by replacing D238 with cysteine for one of the mutations would change the surface charge so much that it would be difficult to assess the influence of the other mutations. Before proceeding with the mutations, we analyzed the behavior of the LCC^ICCG^ (studied further below), LCC^IG^ and PES-H1^FY^ variants in solution through 100 ns MD simulation each. Comparison of their root mean square deviation and fluctuation (RMSD and RMSF) to the corresponding WT enzyme confirms their stability and similar dynamics (Fig. S1).

The mutations we selected mainly involved polar and positively charged residues, which we replaced by apolar and negatively charged residues of similar size. The aim is to increase the overall mobility on the PET surface so that proximal binding becomes more likely. Hence, by replacing positively charged residues, we intend to avoid stabilizing electrostatic interactions with negative charges at the PET chain ends and the negatively charged regions on the PET bulk surface, as can be seen in the EPS of PET in Fig. S2. Moreover, to induce a strong local effect, three mutation sites in close proximity to each other were selected, where preference was given to residues that were in contact with PET in distal areas for at least 75% in one or more of our initial simulations (Fig. 2A). For each enzyme, two triple mutants were designed to change the *E*_int_ ratio of polar and apolar residues (PES-H1^FY^: T11V/S13A/S14A, Q26L/T27V/T28V; LCC^IG^: L66A/S67A/S69A, T211V/S216A/N239L) and a third triple mutant was created with the aim to to specifically influence the enzyme’s EPS (PES-H1^FY^: S13A/R19E/R75E, LCC^IG^: R143E/S145A/R151E). The LCC^IG^ T211V/S216A/N239L mutations are the only ones located near the active site. We have observed a dominant PET residency in this region, which may impede PET progression from this site to the biding cleft. For the modification of the EPS in PES-H1^FY^, it was not possible to identify at least two positively charged residues that i) are in close proximity to each other, ii) were in lasting contact with PET in the simulations of PES-H1 and iii) whose replacement would not affect protein stability due to their involvement in intra-protein salt bridges. Therefore, R75 was included in the corresponding variant, PES-H1^FY^ S13A/R19E/R75E, even though it did not make contact with PET for more than 75% of the simulation time. For LCC^IG^ L66A/S67A/S69A, we deviated from the strategy of mutating polar and positive residues, as L66 appeared to favor PET binding, with more than 75% retention time in two of the six simulations. It should be mentioned that among the hydrophobic residues, leucine was always the strongest mediator for PET interactions. For LCC, L66 had the strongest effect in this regard, and due to its proximity to S67 and S69, we included it in the corresponding mutation.

For each of the LCC^IG^ and PES-H1^FY^ variants, we performed 100 ns MD simulations in triplicate starting from the snapshot of previous simulations where LCC or PES-H1 had the lowest distance between the PET bulk and the three mutation sites.

The simulations were analyzed with respect to *E*_int_ and the possible reorientation of the enzyme at the PET surface. While the latter could not be observed in the simulations, we were able to alter *E*_int_ and also the EPS (Fig. 3). For both enzymes a reduction in their interaction with PET was achieved, i.e., the *E*_int_ values have become less negative. The most significant *E*_int_ changes occurred for the LCC^IG^ variants, which is consistent with our design strategy of making the LCC (IG) surface more similar to that of PES-H1 (FY) to facilitate the reorientation of the enzyme on the PET surface. The introduction of negatively charged residues successfully changed the EPS in this region from positive or neutral to negative and significantly increased *E*_int_ for the corresponding LCC^IG^ mutations (R143E, R151E) and to a lesser extent for the PES-H1^FY^ mutation R19E. The latter can be explained by the fact that the PES-H1^FY^ surface was already rather negative before this mutation, which reduces the mutation effect. PET did not interact with position 75 in any of the simulations with PES-H1^FY^ and PES-H1^FY^ S13A/R19E/R75E. This is probably because the starting structure chosen for the simulations had the residue 75 farthest from PET among the three mutation sites. Therefore, the effect of the R75E mutation cannot be quantified. The implementation of apolar residues, especially alanine, at the expense of polar residues also significantly increased *E*_int_ (e.g., S145A in LCC^IG^, S13A in PES-H1^FY^), while larger apolar residues such as valine or leucine yielded no significant changes or even made the interaction at these sites more favorable (e.g., T211V and N239L in LCC^IG^; Q26L and T28V in PES-H1^FY^). In conclusion, local PET binding was successfully reduced in our simulations by introducing small apolar alanine or negatively charged residues. This is hypothesized to promote the progression of the enzyme–PET contact area towards the binding cleft, thereby enhancing PET binding and hydrolysis.

**Figure 3:**
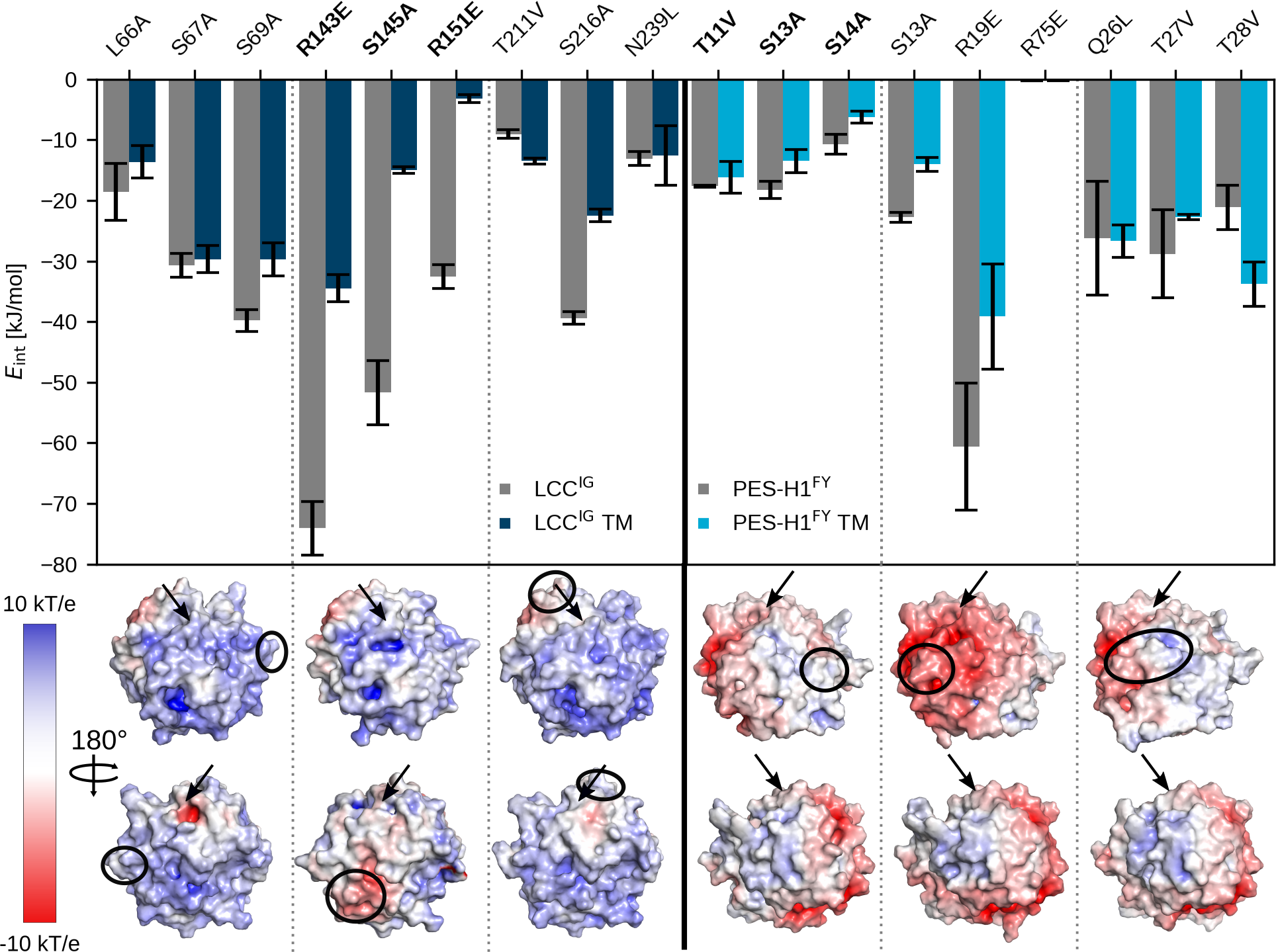
Results for the *in silico* mutation of surface residues for LCC^IG^ (left) and PES-H1^FY^ (right). (Top) The interaction energies (*E*_int_) between PET and the residues at mutation sites are shown for the parent enzyme IG or FY (gray) and the three triple mutants (TMs) per enzyme (dark blue for LCC^IG^, light blue for PES-H1^FY^). The mutations are given at the top, the TMs are separated by vertical dotted lines and the TMs, which were evaluated experimentally are highlighted in bold. The *E*_int_ values are averages over three simulations with the SEM given by error bars. (Bottom) The EPS of the TMs are shown. The locations of the mutations are denoted by black circles and the catalytic site is indicated by an arrow.

#### Distal mutations have a major influence on PET degradation activity

To test our hypotheses derived from the *in silico* studies, two of the triple mutants were subjected to *in vitro* analysis: PES-H1^FY^ T11V/S13A/S14A with increased hydrophobicity and LCC^IG^ R143E/S145A/R151E with increased negative charge. PES-H1^FY^ and the LCC^ICCG^ variant were used as reference enzymes. The latter was used instead of LCC^IG^ to allow for a better comparison to published results. ^3,14^ The PET degradation activity was determined with amorphous PET films under specific reaction conditions favored for each enzyme.^14^ For PES-H1^FY^, 1 M potassium phosphate buffer has ensured the most efficient hydrolysis rates, which was presumably attributed to its stabilizing effect.^14,61,62^ For an optimal comparison with FY, LCC^ICCG^, which is not as salt-dependent as PES-H1^FY^,^3^ was also investigated in 1 M reaction buffer. Unfortunately, the LCC^ICCG^ triple mutant could not be expressed, possibly due to the altered surface charge, which could introduce repulsive forces that interfere with native folding.

The percentage weight loss of the PET films after 24 h was determined to evaluate the enzymatic PET degrading activity. As shown in Fig. 4 the substitution of three residues in PES-H1^FY^ far away from the binding site led to greatly reduced activity. This observation does not fully agree with our hypothesis that decreasing the surface polarity far away from the PET binding site should encourage proximal PET binding and thus increase the PET degrading activity. On the other hand, the *in vitro* results confirmed our hypothesis that the initial binding to the PET surface plays an important role for the enzymatic activity. This binding occurs far away from the active site, ruling out any direct interference with the hydrolysis reaction, as shown with the current PES-H1^FY^ variant T11V/S13A/S14A (Fig. 3). Therefore, future studies should thoroughly investigate mutations on the LCC and PES-H1 surface that enhance PET degradation.

**Figure 4:**
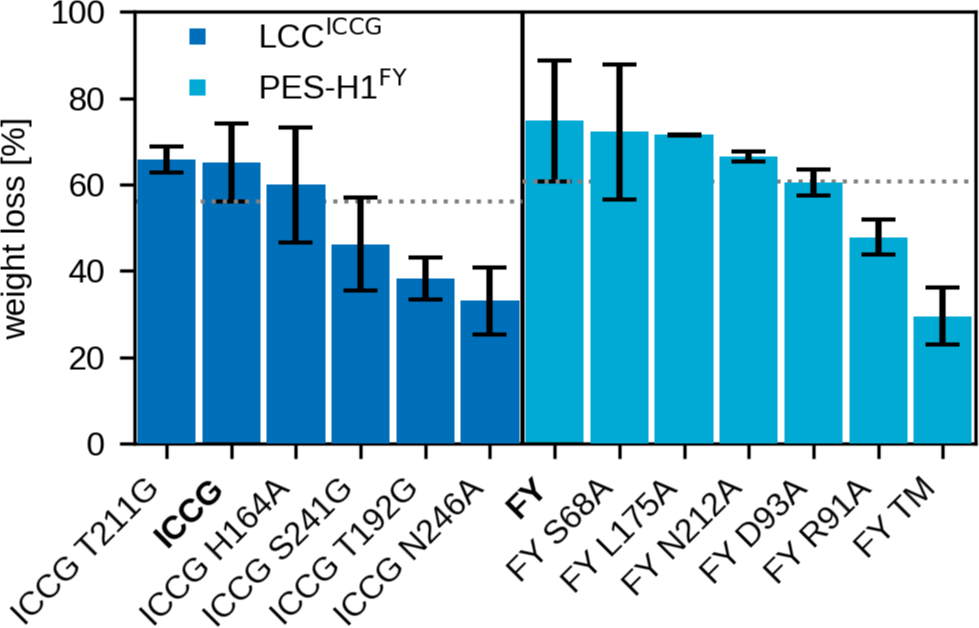
Results from the *in vitro* testing of the original and variants of LCC^ICCG^ (ICCG, left, blue) and PES-H1^FY^ (FY, right, light blue). The variants include five single-point mutations per enzyme, given as axis labels, and the PES-H1^FY^ triple mutant (TM) T11V/S13A/S14A. Variants are ordered according to the percentage weight loss of PET films upon degradation, which serves as a measure for the PET degradation activity. Depolymerization of 60 mg PET film was conducted for 24 h at 70 °C in 1 M potassium phosphate buffer (pH 8) with 60 µg of respective purified enzyme variants. Reactions were performed in duplicate for PES-H1^FY^ L175A and LCC^ICCG^ T211G and for all others at least in triplicate.

### 2.2 Entry of PET into the binding site

#### The energy barrier associated with PET entry can be overcome by enhanced sampling, leading to productive binding poses

Since the entry of PET into the binding cleft was not observed with conventional MD simulations, our hypothesis is that a considerable energy barrier must be overcome for this to occur. Therefore, we performed HREMD simulations to increase the sampling of the conformational space of PET. In these simulations, the energy function of PET is changed in a way that the energy barriers for its conformational changes become smaller. We applied these changes to five of the six replicas we considered per HREMD simulation, with the height of the barriers decreasing with increasing replica number. The changes applied to the energy function correspond to increased simulation temperatures, which we will use in the following discussion. For the starting structures of these HREMD simulations, we extracted three conformations from the previous 2 µs MD simulations, in which the binding of the enzyme to a PET bulk was investigated, with the selection criterion of a distance between the catalytic serine and PET of at most 5 Å. In addition, we retained only the PET-9mer closest to the enzyme and removed four units from it, leaving a PET-5mer closest to the catalytic serine (Fig. 5A). This resulted in different starting structures for both enzymes, which were submitted to HREMD simulations with 115 ns per replica. For LCC we performed five HREMD simulations and four for PES-H1 (Tab. S1). In these simulations, and later in the simulations for LCC^IG^ and PES-H1^FY^, we found that one of these three starting structures (designated 4thL) had the highest number of PET-5mer entry events. Therefore, for the later HREMD simulations of the enzyme variants, this particular starting structure was preferentially used, unless the mutation site was significantly closer to the PET-5mer in another starting structure, which was then used instead. These HREMD simulations enabled to observe several entry events of the PET-5mer into the active site and how this led to the formation of productive binding states, which we distinguish into pro-S and pro-R states (Fig. 5B). A productive state is present when the distances between the carbon of the nearest PET-5mer ester and the γ-oxygen of the catalytic serine (SER–C distance) and between the carbonyl oxygen of the same ester and the two amino hydrogens of the oxyanion hole scaffold are sufficiently small to allow hydrolysis. As distance cutoff we defined *≤*4 Å here. Pro-S and pro-R are defined as PET poses that promote the formation of an S- and R-type tetrahedral intermediate, respectively.^63^ An example of the PET-5mer entry into the binding cleft is shown in Fig. 5C. Here, 12 ns of one of the HREMD simulations of LCC can be seen, where the SER–C distance is initially above 6 Å (the outer phase) and drops to below 4 Å (inner phase) within *≈*2 ns (entry phase).

**Figure 5:**
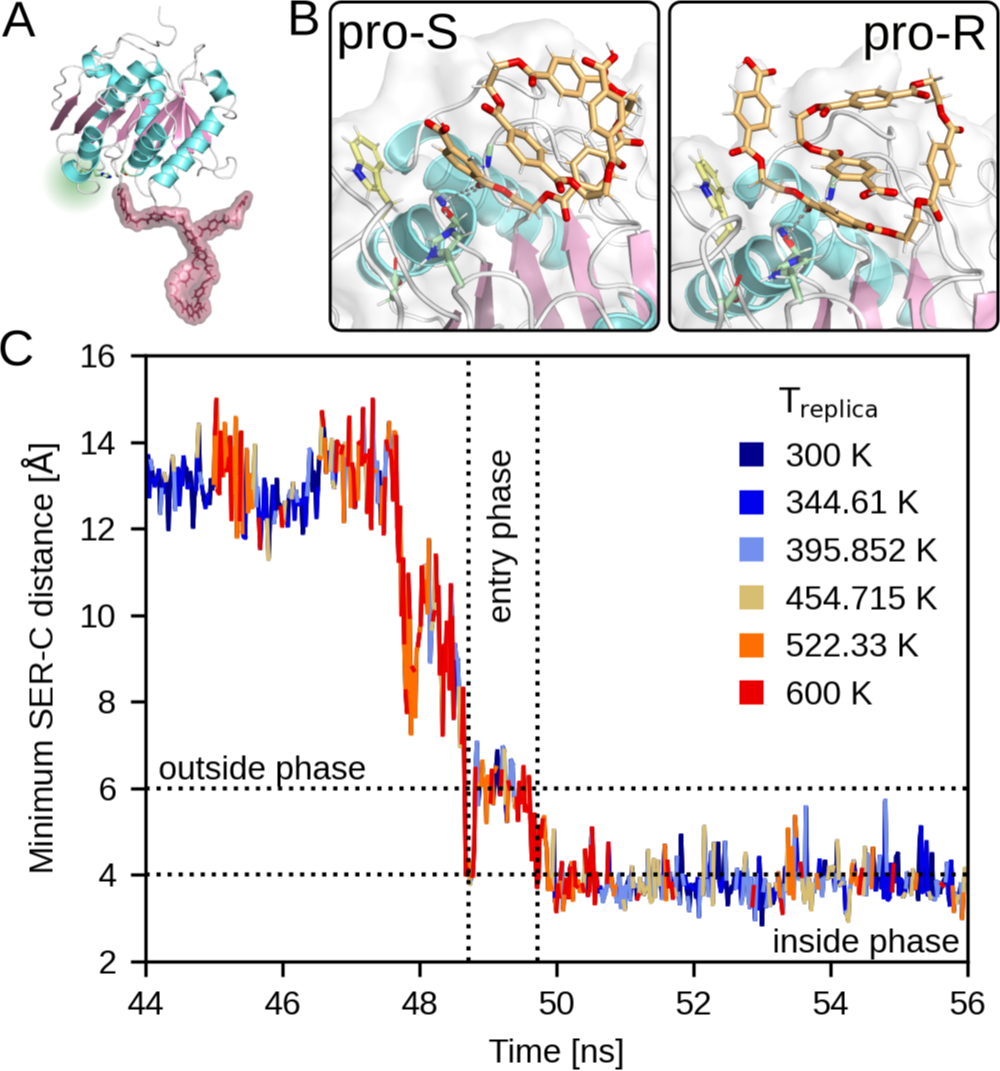
Setup for HREMD simulations and PET-5mer entry results. (A) An exemplary HREMD starting structure is shown to demonstrate the initial distance between the PET-9mer and the enzyme. (B) PET-5mer entry into the active site resulted in pro-S and pro-R poses, for which representative snapshots are shown. (C) The entry of the PET-5mer to the active site was monitored by the evolution of the SER–C distance. Here, a representative entry is shown for LCC. The entry was facilitated by the replicas at high energies/temperatures (colored in orange to red), while conformations from replicas simulated at low energies/temperatures are colored in shades of blue (see color key for exact values). The dotted lines indicate the boundaries for the outside, entry, and inside phases, which are defined by the distance criteria (horizontal lines) and resulting times (vertical lines).

Importantly, entry is enabled by exchanging low-energy replicas (corresponding to low temperatures, shown in blue) for high energy replicas (corresponding to high-temperatures, shown in red) immediately before and during entry. Once the entry is complete, the system remains in low-energy replicas. This observation explains why PET entry did not occur in the earlier conventional MD simulations and confirms our assumption that this process involves a significant energy barrier that must be overcome. Since the current results could be due to the changes we made to the system, i.e., removing the PET bulk, we performed conventional 115 ns MD simulations for the three HREMD starting structures (two LCC and one PES-H1 simulations) to see if PET-5mer entry still occurs. However, this was not the case and no productive PET state were sampled. Additional MD simulations were conducted at 343 K, a temperature recently proposed as optimal for enzymatic PET hydrolysis. ^4,64^ These simulations are crucial for surpassing the energy barrier for PET entry. The simulations resulted in pro-R and pro-S states for PES-H1 and LCC when simulated with a single PET-5mer. One of the LCC simulations that encompassed the entire PET bulk yielded a pro-S state.

In order to decipher the source of this energy barrier, we conducted further analyses and defined specific criteria for the identification of productive PET-5mer entries and their corresponding outside, entry, and inside phases: (1) An entry leading to the adoption of a productive state, either pro-S or pro-R, must have taken place and involve a decrease in the SER–C distance from >6 Å (outside phase) to <4 Å (inside phase). (2) The entry phase must last at least 100 ps and a maximum of 5 ns to minimize noise through in and out movements. (3) The inside and outside phases must each last at least 1 ns. The inside phase ends when the SER–C distance exceeds 6 Å and the outside phase ends when this distance falls below 4 Å, or, in the case of more gradual changes, when the SER–C distance exceeds 4 Å on average over the entire inside phase and falls below 6 Å on average over the entire outer phase. Based on these criteria, we identified 29 productive entries for LCC and 15 for PES-H1, corresponding to 4.35% and 6.49% of the corresponding MD frames. Of these entries, 13 and 7 resulted in pro-S poses for LCC and PES-H1, respectively, while the others led to pro-R poses. The pro-R pose was identified via docking studies performed by Tournier *et al.*,^3^ but others later found in QM/MM studies that the pro-S pose is energetically more favorable^63,65^ and that robust binding of a state unfavorable for hydrolysis could inhibit PET degradation activity. ^66^ Thus, although several pro-R poses were sampled in our study, hydrolysis is expected to occur mainly via pro-S poses. Our analysis of the PET-5mer binding events further shows that the two units that share the ester bond to be hydrolyzed are tightly bound to the enzyme, which is required for the reaction to proceed, while the other PET units interact rather loosely with the enzyme and remain flexible. This finding is evidenced by the different snapshots in Figs. 5B, S3 and S5, which show the conformational freedom of PET units that do not interact directly with the active site. This finding is in contrast to earlier assumptions about a long binding cleft that has several subsites to bind several PET units simultaneously.^46,67^

From the analysis thus far it can be concluded that PET has to overcome an energy barrier for entry into the active site to adopt productive conformations. This energy barrier could be driven by conformational changes of PET to fit into the binding cleft. This is supported by recent studies suggesting that a ‘wrapped’ PET conformation is energetically favored in solution, while a ‘W-shaped’ extended conformation, which is crucial for productive binding poses, is observed in association with the enzyme at elevated temperatures. ^68^ Such conformational changes, and thus the associated energy barrier, could be influenced by PET–PET and residue–PET interactions, which is investigated next by analyzing intramolecular PET interactions and intermolecular PET–enzyme interactions during productive PET entries.

#### Intramolecular PET interactions hinder necessary conformational changes for entry into the binding cleft

For successful PET binding in the active site of the enzyme, a polymer chain or a part of it has to detach from the amorphous or crystalline PET bulk, which means that nonbonded interactions such as hydrogen bonds and π-interactions within the same chain and with other PET chains must be resolved, in addition to meeting sterical requirements. To assess the impact of the intramolecular PET-5mer interactions on the energy barrier during entry to the binding cleft, we calculated *E*_int_ between PET units during productive entries. In several instances, we observed robust interactions between multiple PET units prior to entry, suggesting an inhibitory effect (Fig. S3A). Conversely, individual energy peaks corresponding to favorable intramolecular interactions were detected shortly before or during entry, which probably facilitate the necessary structural rearrangements and interrupt the inhibitory interactions (Fig. S3B). In addition, in some cases, entry-promoting effects were found due to newly formed interactions between two units during and/or after entry (Fig. S3C).

To more comprehensively assess the predominant interactions throughout all simulations of both LCC and PES-H1, as opposed to focusing on individual entry events, we concatenated the HREMD data using all replicas, with the higher replicas weighted, for either enzyme and calculated 2D free energy surfaces (FESs) (Fig. S4). One coordinate in these FESs are the interaction energies between two PET units that are not directly adjacent, which allows us to monitor the intramolecular PET-5mer interactions. To correlate this with the PET-5mer entry, we used the SER–C distance as a second coordinate and distinguish between productive and non-productive regions with a cut-off value of 4 Å. All FESs indicate that non-productive PET-5mer states and also those during entry with SER–C distances below 6 Å but above 4 Å are energetically stabilized, which suggests that PET–PET interactions hinder rather than promote PET-5mer entry into the active site.

In our opinion, the most promising approach to overcome this inhibitory effect is to improve the preprocessing of PET waste. Tarazona *et al.* studied the surface erosion process of PET to differentiate between the PET surface and the bulk properties, which might enable to find a sweet spot between increased PET surface mobility and the increased bulk crystallization.^19^ They found that enlarging the PET surface, e.g. by using micronized PET particles, increasing the surface amorphization and lowering the surface glass transition temperature due to the plasticization effect of water promotes the degradation activity by increasing the PET accessibility and mobility to overcome intramolecular constraints. This is also consistent with a recent study by Schubert *et al.*, in which chain mobility and the number of available sites for endo-type chain scission, which depend on the degree of crystallinity, are postulated as the activity-limiting factor causing the initial lag phase for the release of the soluble product.^7^ Another biotechnological approach to mitigate this hindering effect could be to design enzymes to introduce residues that enhance enzyme–PET interactions, which would promote and counterbalance the loss of PET–PET interactions upon entry into the binding cleft.

#### PET–enzyme interactions can also hinder PET entry

Next, we analyzed the contribution of intermolecular interactions between the PET-5mer and enzyme to the observed overall energy barrier during PET-5mer entry. To this end, we first examined individual entry histories and identified effects that promote or inhibit entry by calculating *E*_int_ between the PET-5mer and residues within 3 Å during entry. PET-5mer entry can be hindered by strong PET–enzyme interactions prior to entry or by repulsion of PET-5mer within the binding cleft, so that PET-5mer states outside the cleft are favored (Fig. S5A). Conversely, an entry-promoting effect has favorable interactions during entry, which facilitate structural PET-5mer rearrangements that overcome PET–PET interactions (Fig. S5B). Furthermore, PET-5mer entry gets supported through stabilizing interactions for productive PET5-mer poses, which compensate for the loss of PET–PET interactions (Fig. S5C).

To obtain a more general overview, FESs were created using a similar approach as before, but this time for the PET–enzyme interactions. The SER–C distance was again used as a coordinate to monitor the entry process, while *E*_int_ between the PET-5mer and selected residues was used as another coordinate. The selection and mutation of certain residues allow us to investigate the specific, and sometimes different effects of amino acids in both enzymes. For instance, depletion of tyrosine in the LCC^ICCG^ variant (Y127G) increased activity compared to WT-LCC, while introduction of tyrosine in the PES-H1^FY^ variant (Q94Y) at the corresponding position resulted in increased activity compared to WT-PES-H1.^3,14^ Two more positions are relevant for the improved activity of LCC^ICCG^ and PES-H1^FY^: the L92F mutation in PES-H1^FY^ and the F243I mutation in LCC^ICCG^.^3,14^ The F125L mutation in the LCC, corresponding to position L92F of PES-H1^FY^, has been shown to reduce LCC binding to PET surfaces, increasing its availability for targeting degradation intermediates and enhancing overall activity, so it is also analyzed here.^60^ Additionally, we studied L209 of PES-H1 corresponding to position F243I of LCC^ICCG^, for which a mutation to alanine was chosen instead of isoleucine based on studies reporting alanine as the only variant exhibiting activity close to that of the WT.^14,32^

To investigate the effects of these mutations, we performed HREMD simulations for the LCC^IG^ and PES-H1^FY^ variants as well as for LCC^IG^ F125L and PES-H1^FY^ L209A, and calculated the FES from the HREMD data as described above. The resulting FESs reveals distinctive PET–enzyme interaction patterns for WT and mutated residues (Fig. 6). Mutations at positions PES-H1^FY^ 209/LCC^IG^ 243 stabilize productive states in both enzymes, which can be inferred from new free energy minima (in yellow/brown colors in Fig. 6) at SER–C distances below 4 Å and less pronounced minima for distances above 6 Å after mutation, suggesting weaker interactions with the PET-5mer in latter states. In contrast, the effects at positions PES-H1^FY^ 92/LCC^IG^ 125 and PES-H1^FY^ 94/LCC^IG^ 127 differ between the two enzymes. In the case of LCC^IG^, entry of the PET-5mer after either mutation is promoted due to the elimination of non-productive energy minima resulting from weakened interaction with the PET-5mer for SER–C distances above 6 Å, whereas in PES-H1^FY^, the entry of the PET-5mer after the mutation becomes more difficult due to increased interaction with the PET-5mer in non-productive states. We therefore conclude that the increase in PET degradation activity could result from a direct promotion of PET entry by the mutated residues for LCC^ICCG^, but from different mechanism for PES-H1^FY^.

**Figure 6:**
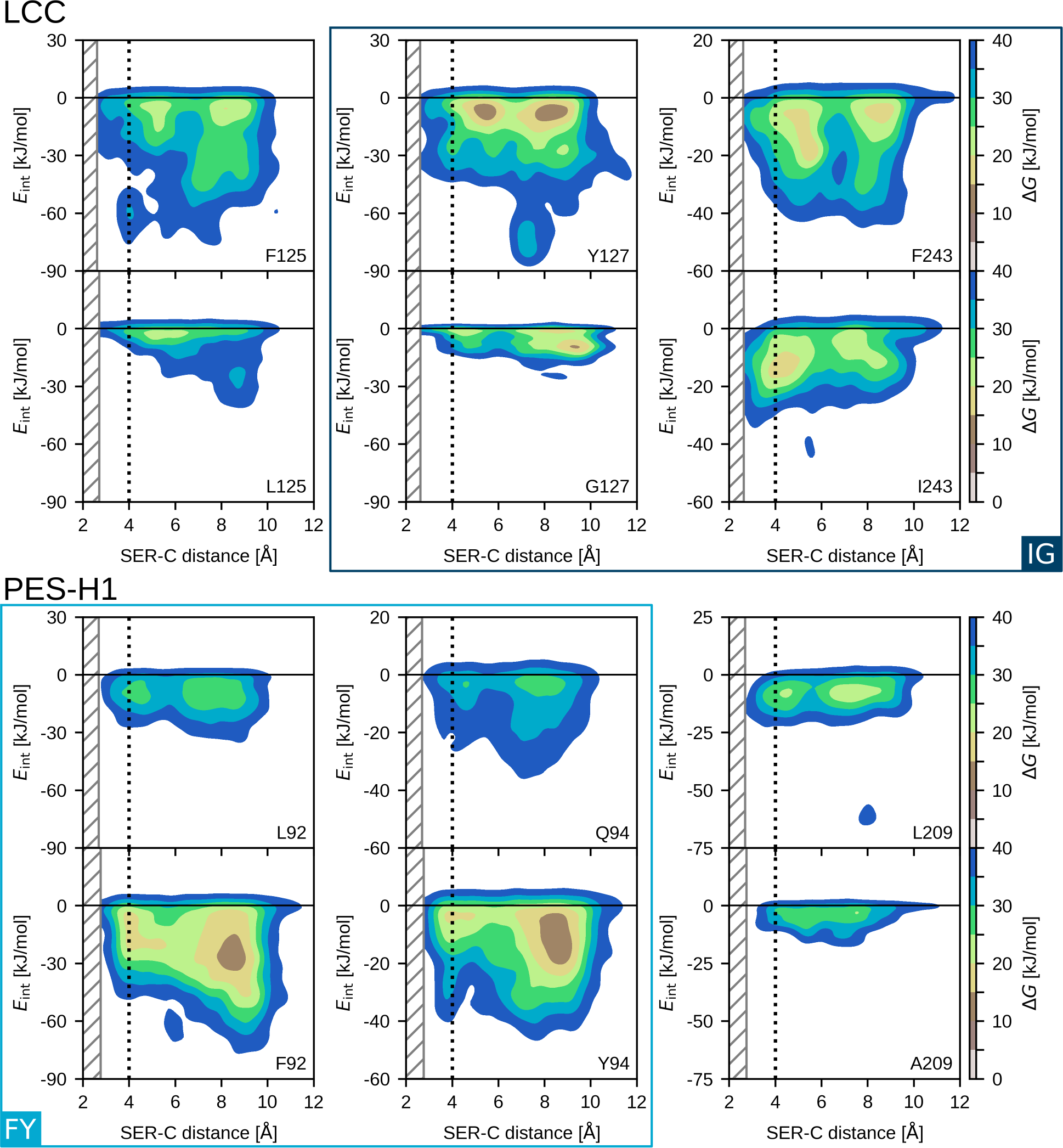
Free energy surfaces of the PET–residue interactions over the SER–C distance obtained from HREMD simulations of LCC and PES-H1 variants. The top row per enzyme presents the FESs for residues present in the WT and the bottom row for the mutated residues in the LCC^IG^/PES-H1^FY^ variants (indicated by blue boxes) and for the LCC^IG^ F125L/PES-H1^FY^ L209A variants. The area for SER–C distance *<* 4 Å harbors productive poses. The gray hatched area indicates distances that were not sampled as the ester carbon atom would be too close to the catalytic serine. The free energies, Δ*G* are given in kJ/mol according to the color scale on the right.

To further explore our hypothesis that increased entry of PET into the active site contributes to increased activity as presumed for LCC^ICCG^, we analyzed additional residues near the active site of both enzymes and then mutated them. Our aim was to identify residues that lack entry-promoting FES features and are therefore promising mutation targets to improve PET entry ability. FES features to be introduced include the creation of free energy minima in the productive region with a SER–C distance below 4 Å, indicating stabilized productive positions of PET-5mer, the elimination of free energy minima in the non-productive region, indicating reduced interactions with PET-5mer, and the facilitation of the transition from the non-productive to the productive region by avoiding energy barriers there. The residues that we identified as promising mutation sites were then mutated to glycine or alanine; larger amino acids were not introduced to avoid creating steric hindrances. In addition, the PES-H1^FY^ H184S variant was simulated, in which the mutation site is located near a highly conserved tryptophan (PES-H1: W155, LCC: W190) adjacent to the binding cleft and whose ‘wobbling’ presumably increases catalytic activity. ^45^ Mutation of H184 to a serine is likely to increase the flexibility of this tryptophan due to the lower steric requirements of serine.

This resulted in a total of 14 mutation sites that were evaluated for each enzyme. For each mutant, an HREMD simulation was performed (see Table S1 for the list of HREMD simulations) and the FES calculated as described above. In many cases the overall goal of the *in silico* mutations to facilitate PET-5mer entry into the active site was achieved. For these cases the FES is shown in Fig. S7, and they include mutations with improved stabilization of productive states (PES-H1^FY^: S68A, D93A; LCC^IG^: T211G, N246A), reduced interaction in non-productive states (PES-H1^FY^: S68A, R91A, D93A, N212A; LCC^IG^: T192G, S241G), or smoother transitions towards productive states without intermediate states where the PET-5mer could get trapped (PES-H1^FY^: S68A, D93A; LCC^IG^: HIS164A, T211G, S241G, N246A). The PES-H1^FY^ H184S variant could neither improve the interactions at position 184 nor the interactions of the ‘wobbling’ tryptophan with the PET-5mer. The five most promising *in silico* mutations per enzyme (i.e., the ones shown in Fig. S7) were then evaluated experimentally, using the same protocol as employed for the previous PES-H1^FY^ triple mutant to determine the PET degradation activity. It should be noted that one of these mutants, PES-H1^FY^ L175A, is a borderline case, as a state slightly above the 4 Å criterion applied to the SER–C distance became stabilized; we nonetheless included it in our experimental evaluation, which is discussed below.

#### Kinetic experiments reveal an increase in activity for the PES-H1^FY^ S68A mutation

In order to link simulation and experiment, the activity of the mentioned mutations introduced in the PES-H1^FY^ and LCC^ICCG^ enzymes was determined. The percentage weight loss of the PET films shown in Fig. 4 indicates that none of the variants resulted in a significant increase in the degradation activity. PES-H1^FY^ R91A, LCC^ICCG^ T192G, and LCC^ICCG^ N246A showed significantly lower degradation activity than PES-H1^FY^ and LCC^ICCG^, respectively. Interestingly, these are also the three variants with the least promising FES profiles among the ten selected variants (Fig. S7). The FES improvements in these mutants were mainly related to the destabilization of the non-productive states, while the productive state did not gain in stability. In LCC^IG^ N246A, on the other hand, while the productive state was stabilized, also a non-productive state at SER–C distances of *≈*6.5 Å became more stable, which could explain the reduction in activity for this variant. In addition, there is also the possibility that other factors may influence the experimental outcome and thus mask improvements due to mutations. It should also be mentioned that the reproducibility of the experiments was problematic despite uniform experimental conditions in all experiments, which we suggest can be attributed to influences of the inherent properties of the PET films such as crystallinity or molecular weight distribution. ^16^

To reduce potential masking effects and identify subtle effects on individual steps in PET degradation, we analyzed the degradation kinetics using a previously described turbidimetric assay using PET nanoparticles (PET-NP) that measures turbidity changes at 600 nm over time.^14,69^ Here, not only the substrate load [*S*_0_] was varied while maintaining a fixed enzyme concentration [*E*_0_], as done in conventional Michaelis–Menten kinetics measurements (^conv^MM), but also the reverse approach with varying enzyme concentration and fixed substrate load was followed, which is called inverse Michaelis–Menten (^inv^MM) and is a commonly used approach for kinetic measurements with insoluble substrates such as PET.^20,29,70–76^ To further ensure a more homogeneous distribution of the substrate, PETNP were used instead of a PET film, which should lead to more uniform and reproducible results.^76^ This approach was used to evaluate the three most interesting candidates per enzyme that showed the most improvement in their FES profiles upon mutation while exhibiting similar PET degradation activity to PES-H1^FY^ and LCC^ICCG^, respectively, namely PES-H1^FY^ S68A, D93A, N212A and LCC^ICCG^ H164A, T211G, S241G (Tab. 1 and Tab. S3).

**Table 1:** The Michaelis–Menten constants *K*_m_ were derived from the nonlinear fit to experimental data for both settings, ^conv^MM and ^inv^MM, and both enzymes, LCC^ICCG^ (ICCG) and PES-H1^FY^ (FY), and corresponding variants. Additional secondary parameters *k*_cat_, ^molar^*K*_m_ and ^molar^*η* are provided.

| | $^{\text{conv}}K_m \left[ \frac{\text{g}}{\text{L}} \right]$ | $^{\text{inv}}K_m [\mu\text{M}]$ | $k_{\text{cat}} \left[ \frac{1}{\text{min} \cdot \mu\text{M}} \right]$ | $^{\text{molar}}K_M [\mu\text{M}]$ | $^{\text{molar}}\eta \left[ \frac{1}{\text{min} \cdot \mu\text{M}^2} \right]$ |
| --- | --- | --- | --- | --- | --- |
| LCC<sup>ICCG</sup> | 0.215 ± 0.059 | 0.365 ± 0.061 | 0.047 | 0.357 | 0.132 |
| ICCG H164A | 0.095 ± 0.063 | 0.861 ± 0.164 | 0.032 | 0.278 | 0.114 |
| ICCG T211G | 0.091 ± 0.068 | 0.430 ± 0.130 | 0.042 | 0.223 | 0.189 |
| ICCG S241G | 0.250 ± 0.068 | 0.234 ± 0.070 | 0.054 | 0.357 | 0.151 |
| PES-H1<sup>FY</sup> | 0.391 ± 0.109 | 0.892 ± 0.105 | 0.088 | 1.036 | 0.085 |
| FY S68A | 0.357 ± 0.117 | 0.278 ± 0.053 | 0.084 | 0.728 | 0.115 |
| FY D93A | 0.501 ± 0.180 | 2.017 ± 0.481 | 0.040 | 2.434 | 0.016 |
| FY N212A | 0.481 ± 0.174 | 1.767 ± 0.318 | 0.068 | 1.564 | 0.043 |

It should be noted that the conventional measurements led to inconsistent results, especially for the LCC^ICCG^ variants, which confirms that the conventional approach and the applied kinetic model meets experimental challenges when dealing with PET as substrate. Therefore, the following discussion is based on results obtained with the inverse Michaelis–Menten approach. For ^inv^*K*_m_ significant differences are observed in enzyme–substrate affinity between some of the variants and the respective parent enzyme, LCC^ICCG^ or PES-H1^FY^. An approximate 3-fold decrease in ^inv^*K*_m_ was detected for the PES-H1^FY^ S68A variant compared to PES-H1^FY^ (PES-H1^FY^ S68A: ^inv^*K*_m_=0.273 µM; PES-H1^FY^: ^inv^*K*_m_=0.892 µM), which indicates a higher enzyme–substrate affinity for PES-H1^FY^ S68A. For the other two PES-H1^FY^ variants, higher ^inv^*K*_m_ values than for PES-H1^FY^ suggest a decreased affinity, which is supported by similar trends in the ^molar^*K*_m_ values. For LCC^ICCG^ the results are less clear. The higher ^inv^*K*_m_ values for LCC^ICCG^ H164A and LCC^ICCG^ T211G than for LCC^ICCG^ are not mirrored by the corresponding ^molar^*K*_m_ results, which are lower, while the turnover rate constant *k*_cat_ could also not be improved. For the LCC^ICCG^ S241G variant a reduction in ^molar^*K*_m_, a similar ^molar^*K*_m_ value as for LCC^ICCG^ and an increase in *k*_cat_ were observed, but these improvements are not statistically significant (Tab. S3). Combining affinity and turnover rate into ^molar^*η* = *k*_cat_*/*^molar^*K*_m_ yields the molar catalytic efficiency. First, it should be noted that a higher catalytic efficiency was observed for all LCC^ICCG^ variants than for the FY variants. A comparison of these figures per enzyme indicates that PES-H1^FY^ S68A and LCC^ICCG^ T211G outperform FY and LCC^ICCG^, respectively, while LCC^ICCG^ S241G shows a slight improvement. Thus, the most promising mutations revealed by experiments are also the mutations with the most promising FES changes (Fig. S7). In particular for the T211G mutation in LCC^ICCG^ we succeeded in creating a smooth free energy path leading to the productive state, which corresponds to the largest increase in ^molar^*η* with respect to the parent enzyme for all variants studied. With the S68A mutation in PES-H1^FY^ we were able to abolish the stabilization of unproductive states, while the productive state gained stability, yet not as much as with T211G in LCC^ICCG^.

In our *in silico* mutation strategy, we have restricted ourselves to alanine and glycine as target amino acids. Given the promising, yet not perfect results for the PES-H1^FY^ S68A variant, we decided to investigate the effects of side-chain size, polarity, charge and aromaticity at that position. To this end, we generated additional *in silico* variants (S68I, S68F, S68Q, S68D, S68K), run HREMD simulations per variant and analyzed the resulting FES (Fig. S8). The results reveal that our original strategy to mutate PES-H1^FY^ S68 to alanine was a good choice, as mutating this residue to any other residue stabilizes non-productive states and isolated areas emerged in the FES, suggesting that the transition to the productive state to has become more difficult. We therefore conclude that interactions with this residue are generally not beneficial for PET entry, which is best avoided by removing steric hindrances as realized in the PES-H1^FY^ S68A mutant.

In conclusion, we were able to define PET entry features by comparing the FES of residues of the respective parent enzyme with their mutations, which allowed us to link the FES appearance with the enzymatic activity measured in experiments. The FES approach allowed us to identify residues that potentially hinder PET entry, and PES-H1^FY^ and LCC^IG^ variants were generated in which these residues were mutated and evaluated computationally and experimentally. This analysis confirmed our idea that changing the FES so that PET can more easily enter the active site by removing energy traps before reaching the productive state and stabilizing the latter, which was best realized in PES-H1^FY^ S68A and LCC^ICCG^ T211G and resulted in higher catalytic activity. With respect to the experimental results it should be noted that ^inv^*K*_M_, in contrast to ^conv^*K*_M_, reflects how effectively the enzyme can operate at different enzyme concentrations for a given substrate load. Considering the results of both approaches, it seems conceivable that the efficiency of the enzymes has increased especially at low enzyme concentrations, while their PET affinity has not necessarily improved. This would be advantageous in industrial applications where the substrate concentration is high and the enzyme concentration is relatively low and adaptation is impractical for cost or feasibility reasons.

#### Principal component analysis reveals that PET can enter via three pathways

To complete our comprehensive analysis of the PET entry into the binding cleft, we next present the complete pathway of how the PET-5mer reaches its final productive state in the active site. In doing so, we do not limit our analysis to the assumption that there can only be one such pathway, but attempt to identify all possible pathways. To this end, we compiled data of all entry events with phases inside the active site of 1 ns. Here, we included data from all simulated variants per enzyme, LCC and PES-H1, assuming that the general pathways should not be affected by the mutations, but only the energies along the pathways (as shown in our previous FES analysis), which could shift the balance between pathway preferences in the case of multiple entry pathways. This generated a large dataset that is suitable for a meaningful principal component analysis (PCA). The PCA was applied to the distances of the PET-5mer ester carbon, which becomes productively bound after entry, to all C_α_ atoms of the corresponding enzyme within 10 Å. This allows to capture the dynamics during the PET-5mer entry without introducing bias by pre-selecting specific residues. The resulting principal components (PCs) are used to calculate the free energy (Δ*G*) along the first two PCs, which account for 86% of the variance and are shown as FESs in Fig. S9A,B.

To identify productive and non-productive states in these FES presentations, we projected them into the PC1–PC2 space (indicated by green and rose color in Fig. S9A,B). This shows that the productive states are clearly separated from non-productive states and that the productive states are localized in the Δ*G* minima regions. We further analyzed the contributions of amino acids mainly involved in the PET-5mer entry by their contributions to PC1, which reveals 49 relevant amino acid positions, which account for 62% of the data variance (Fig. S9C). The involvement of so many positions highlights how complicated the entry pathway of a polymer such as PET into the active enzyme site is. This complexity also challenges the applicability of collective variable-based methods, such as metadynamics, ^77^ for the targeted sampling of entry pathways. HREMD simulations, as used in this study, do not restrict the conformational PET space to be sampled, but simply increase the space that is generally sampled, thus providing a suitable approach. Inspection of the FES along the PCs of the enzyme–PET contacts also shows that the PET-5mer does not enter the active site via an exclusive pathway, but has three such preferential entry routes in both enzymes. For each of these three pathways, a representative trajectory is projected onto the FES and also structures of both enzymes (Fig. 7).

**Figure 7:**
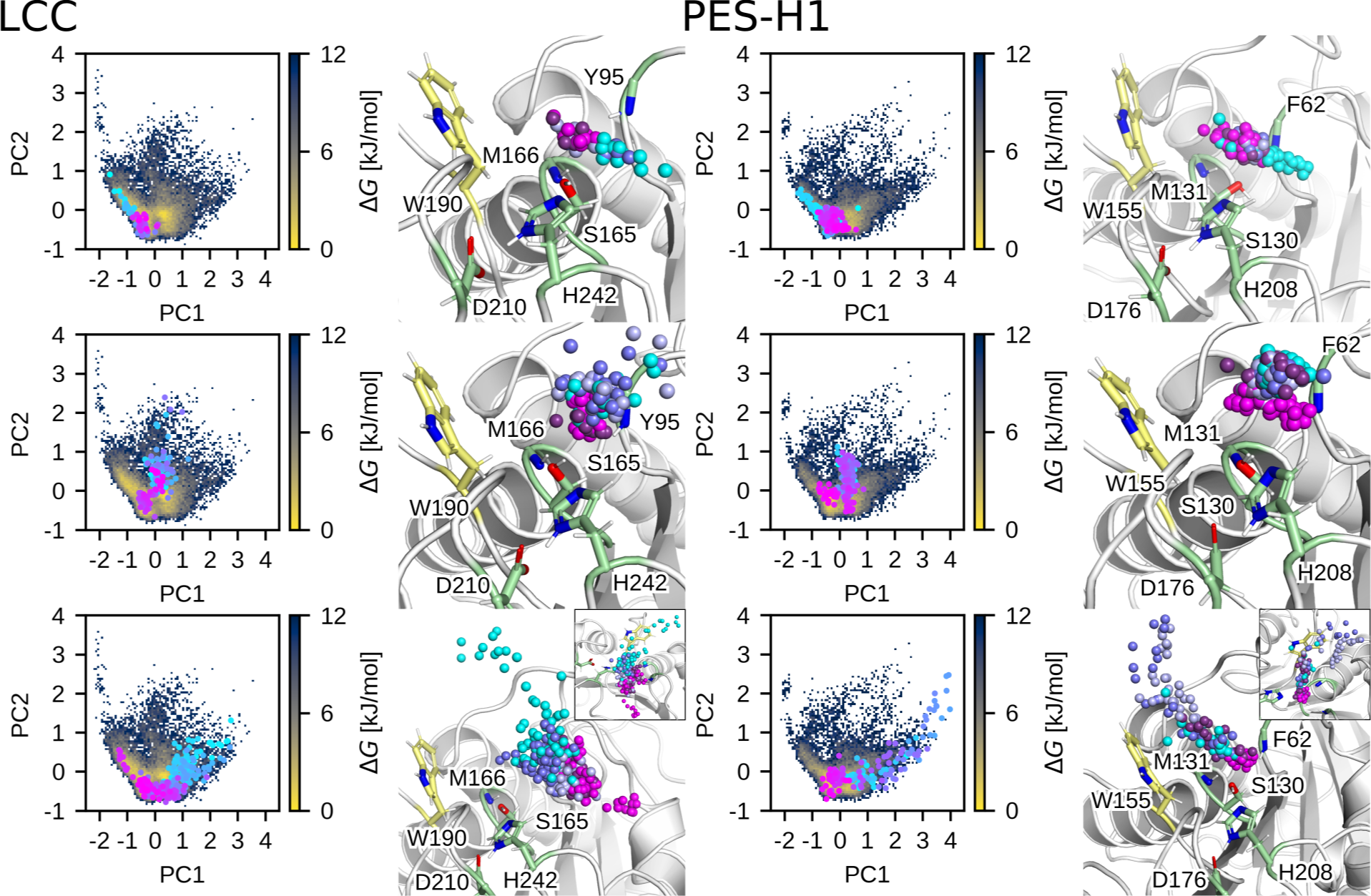
The three preferred entry pathways of the PET-5mer as revealed by PCA applied to the HREMD data involving the simulations of all LCC (left) and PES-H1 (right) variants performed in this work. For each pathway, a representative example is shown. The pathways are mapped as circles on the FES in the PC1,PC2 plane of either enzyme and are also shown (as spheres) together with the corresponding enzyme structure. The progress of each pathway is highlighted by the color change from cyan to magenta. In the structure plots, the spheres show the positions of the PET-5mer ester carbons that are finally productively bound.

These trajectories show how the position of the PET ester carbon evolves towards its productive pose, represented by the color change from cyan to magenta. The first pathway leads from the side opposite the ‘wobbling’ tryptophan to the active site (Fig. 7, top). The second pathway goes directly from the exterior to the catalytic serine (Fig. 7, middle), while in the third pathway, the ester bond slides over the benzene ring of the ‘wobbling’ tryptophan until it reaches its final position for hydrolysis (Fig. 7, bottom). Visual inspection of a number of entry pathways revealed a significant flexibility of the ester carbon position, involving multiple reorientations of the ester position, even when close to the active site, until a final productive conformation is attained. Both enzymes exhibit the same entry pathways, with the first route being the most clearly defined and energetically favored in both cases. As for the second-preferred route, the LCC shows a preference for the direct entry of PET-5mer as found in the second pathway, which is also better defined than in PES-H1, while in the latter these arguments apply to the third pathway.

The pathway preference may be influenced by the interactions with certain residues and their contribution to PC1, the principal component that mainly distinguishes the three pathways from each other (Fig. S9C). Amino acids with a negative contribution predominantly influence the negative range of PC1, impacting the first pathway located at negative PC1 values. Conversely, amino acids with a positive contribution primarily affect the positive range of PC1, influencing the third pathway situated at positive PC1 values. The magnitude of the amino acid contributions reflects their respective strengths. The second pathway corresponds to PC1 *≈* 0 and therefore has no significant amino acid contributions along PC1. The positions mutated in PES-H1 and LCC correspond to amino acids that contribute to PC1. Among the three positions mutated in the more active IG and FY variants, two contribute negatively to PC1 (PES-H1^FY^: 94/LCC^ICCG^: 127 and PES-H1^FY^: 92) and thus primarily affect the first pathway, which has been identified as the most important for both enzymes. The other mutation in LCC^ICCG^ contributes to the positive PC1 direction and consequently to the third pathway (LCC^ICCG^: 243). For several of the positions that we previously analyzed in terms of the FES reflecting their interaction with the PET-5mer, we here find a strong contribution to PC1. In particular, position 68 of PES-H1 makes the largest contribution to PC1, which turned out to be the PES-H1^FY^ variant with the highest catalytic efficiency after its mutation from serine to alanine (Tab. 1). Together with position 209 in PES-H1 (which corresponds to position 243 in LCC and emerged as important for the LCC^ICCG^’s activity^3^), position 68 lines one end of the catalytic binding cleft and might, thus, constitute a sterical hindrance when the predominant entry pathway is attempted (Fig. S6). For LCC, we had identified position 241 as a promising mutation site, which also contributes to the first entry pathway.

In summary, the PET entry process exhibits a certain degree of variability due to the considerable flexibility of the PET chain and persistent PET–PET interactions during entry as shown above, but also due to the possibility that the ester bond that is eventually cleaved can reach the active site by different routes. For both PES-H1 and LCC, we have identified one pathway as the predominant one, which is identical for both enzymes and is consistent with the findings by Falkenstein *et al.*, who proposed this pathway as the PET entry pathway.^52^ However, the plausibility of stepwise PET degradation through ‘hopping and sliding’ and successive cleavage of PET ester bonds as the primary degradation mechanism appears less tenable.

## 3 Conclusion

In this study, we systematically investigated the binding trajectory of PET to LCC and PES-H1, aiming to deepen our understanding of the adsorption process and PET entry into the binding site. To this end, we designed the study approach summarized in Fig. 8. We started with examining the initial adsorption of the PET hydrolases at a large amorphous PET bulk using conventional atomistic MD simulations and explored the subsequent entry of PET into the catalytically active site using enhanced-sampling HREMD simulations. Each simulation step involved analyzing the energetics of the different binding phenomena to identify enzyme residues that either promoted or hindered these events. This analysis was complemented by amino acid mutations, initially evaluated through additional simulations and subsequently validated through experimental measurements in the wet lab.

**Figure 8:**
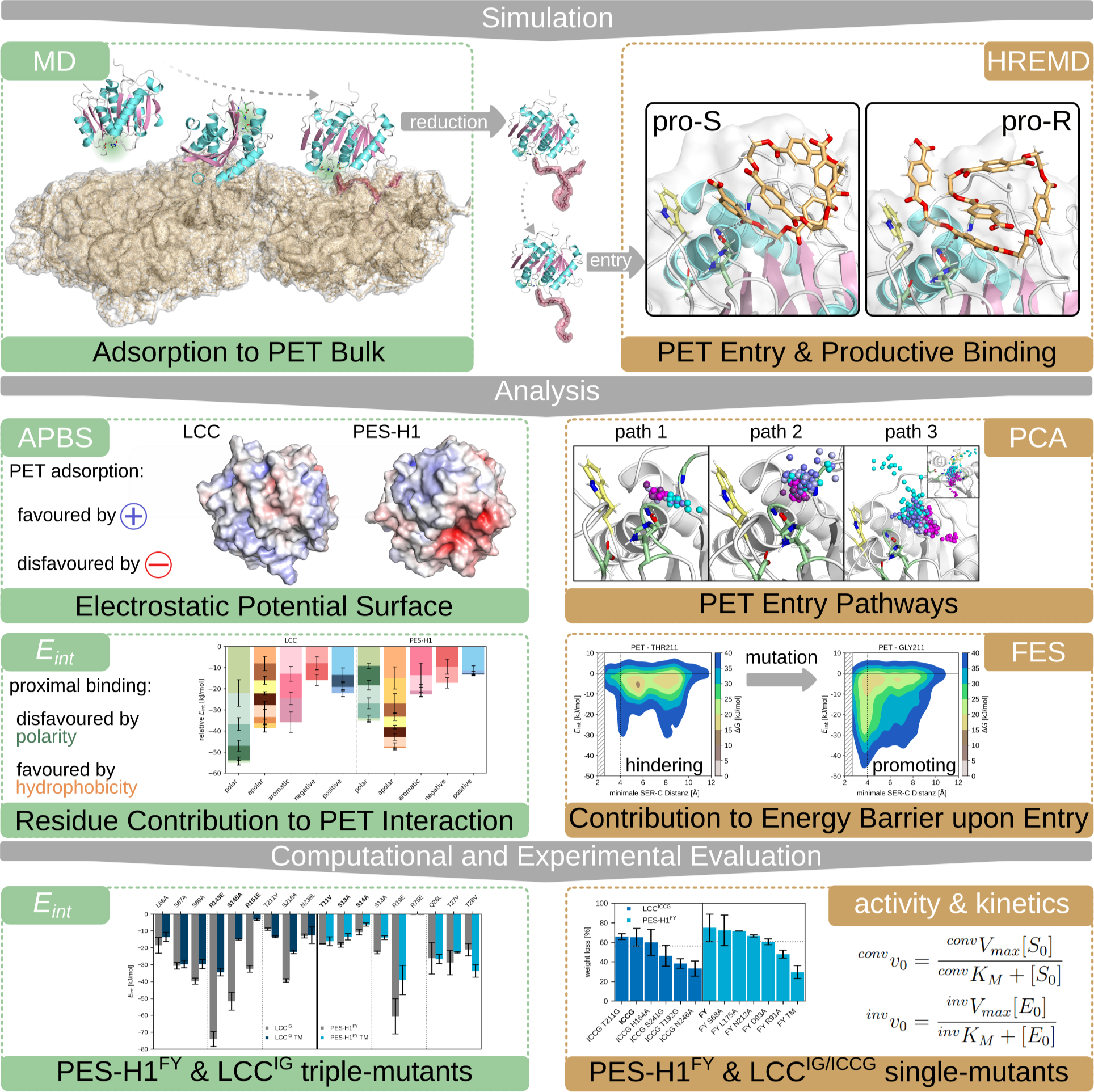
Summary of the workflow applied and results obtained in this study.

Our primary finding regarding the PET surface binding is that heightened adsorption affinity, facilitated by the introduction of positive surface patches and polar residues, may not correlate with improved enzymatic activity. This conclusion aligns with the Sabatier principle, underscoring the importance of balancing binding affinity across enzyme regions, both proximal and distal to the active site, with enzymatic activity optimization. For the next step, the entry of PET into the binding site, we have made the important observation that there is not just one such pathway, but three main ones. In each of these pathways, two critical sources of energy barriers may impede PET from accessing the active site and assuming a productive state. One arises from intra- and intermolecular PET-PET interactions, requiring resolution for the linear binding cleft to accommodate a long, flexible PET chain. The other potential hindrance stems from PET interactions with amino acid residues along the pathway. Our analysis, including free energy surface and principal component analysis of these interactions for wild-type and mutant LCC and PES-H1 enzymes, reveals that energy barriers may result from the stabilization of unproductive states. Conversely, facilitating a seamless energetic transition from unproductive to productive states and stabilizing the latter could enhance enzymatic activity. This theoretical deduction finds support in kinetic experiments involving the S68A mutant of PES-H1^FY^ and the T211G mutant of LCC^ICCG^. However, it should be noted that the variants we identified to facilitate PET incorporation did not show a significant increase in PET degradation activity. This can be explained by the fact that we did not investigate the PET hydrolysis step itself and the subsequent product release, which are also of great importance for enzyme activity in addition to PET entry into the active site. Future studies should therefore investigate the effects of the mutations investigated here on the activation energy barrier of the hydrolysis reaction using QM/MM simulations and their influences on the resulting product complexes.

In summary, our pioneering methodology synergizes HREMD simulations and free energy analyses to dissect substrate entry pathways and pinpoint barriers. This approach unveils critical insights into the intricate pathways traversed by polymers during degradation, enabling the development of mutation strategies to enhance enzyme kinetics. Here, our computational simulations have significantly advanced the understanding of PET hydrolase mechanisms and offer invaluable guidance for future engineering endeavors aimed at maximizing PET hydrolase applicability. Moreover, this approach holds promise for application to a wide range of enzymes beyond PET hydrolases.

## Acknowledgement

A.J. thanks the German Federal Environmental Foundation (Deutsche Bundesstiftung Umwelt) for the financial support by providing a PhD fellowship. A.J is furthermore grateful to her colleague Lara Scharbert for valuable suggestions in figure design. A.J. and B.S. gratefully acknowledge the computing time granted through JARA-HPC (project PETaseMD) on the supercomputer JURECA at Forschungszentrum Jülich, ^78^ the hybrid computer cluster purchased from funding by the Deutsche Forschungsgemeinschaft (DFG, German Research Foundation) project number INST 208/704-1 FUGG, the Center for Information and Media Technology at Heinrich Heine University Düsseldorf, and the Regional Computing Center of the University of Cologne (RRZK) for providing computing time on the DFG-funded (Funding number: INST 216/512/1FUGG) High Performance Computing (HPC) system CHEOPS. R.W. and U.T.B. acknowledge the funding received from the European Union’s Horizon 2020 research and innovation program under grant agreement no. 870294 for the MIX-UP project and no. 953214 for the upPE-T project.

## Supporting Information Available

The following files are available free of charge.

- Material and Methods Section
- RMSD and RMSF of PES-H1, PES-H1^FY^, LCC, LCC^ICCG^ and LCC^IG^
- EPS of PET
- exemplary PET–PET interaction plots during PET entry
- FES plots for PET–PET interaction for WT enzymes
- exemplary PET–residue interaction plots during PET entry
- Mutation sites highlighted in LCC and PES-H1 structures
- FES plots for PET–residue interaction for variants analyzed experimentally and additional PES-H1^FY^ S68 variants
- PC1 contribution profile and PC1/PC2 FES with projections of (non-)productive states
- additional kinetic parameters

## TOC Graphic

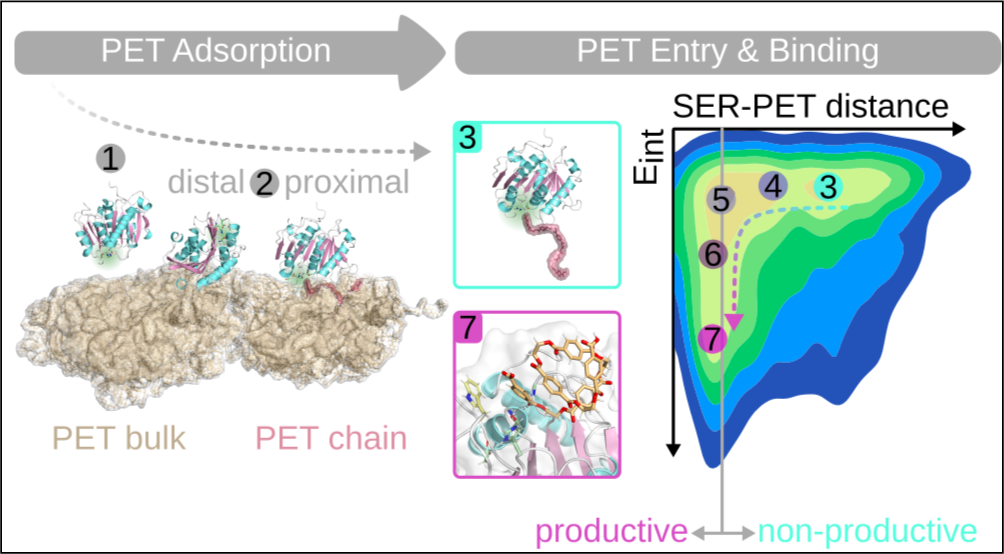

## Notes

### Competing Interest Statement

The authors have declared no competing interest.

