## Supplementary Information for "From Bulk to Binding: Decoding the Entry of PET into Hydrolase Binding Pockets"

#### 1 Methods

##### 1.1 Parameterisation of PET

Quantum mechanics calculations were performed at the HF/6-31G* level for a small PET analogue consisting of three units using Gaussian 09[1] to develop generalized AMBER force field (GAFF)[2] parameters for the terminal and central units. Restrained electrostatic potential (RESP) calculations[3,4] were performed using the antechamber tool[5,6] as available in AmberTools 21[7] to acquire the partial charges, which were redistributed using prepgen to obtain a zero net charge for each PET unit. The input files for the simulation software GROMACS[8] were retrieved using the ACPYPE tool[9] and were already published by Pfaff *et al.*[10]

#### 1.2 MD simulations and analysis protocol

The following protocol was applied for all MD simulations performed in this study, using the software GROMACS version 2020.5.[8] The enzyme and/or PET was placed in a periodic dodecahedron box with at least 10 Å distance to the box edges, solvated using the TIP3P water model[11] and neutralized by addition of the respective amount of sodium or chloride ions matching a physiological concentration of 150 mM. Energy minimization was performed using the steepest descent algorithm[12] until the maximum force fell below 1,000 $kJ\cdot mol^{-1}\cdot nm^{-1}$. The solvent was relaxed in a two-step equilibration approach applying position restraints to enzyme and/or PET heavy atoms: First, a simulation in the NVT ensemble (i.e., with a constant number of atoms (N), constant volume (V), and constant temperature (T)) was followed by a second simulation in the NpT ensemble (i.e., with varying volume, but constant pressure (p)) to set the temperature at 303 K (30°C, v-rescale thermostat[13]) and the pressure at 1.0 bar using isotropic pressure scaling (Berendsen barostat[14]) before subjecting the system to the final production run in the NpT ensemble (303 K, Nosé-Hoover thermostat;[15,16] 1.0 bar, Parrinello-Rahman barostat[17]). During both, equilibration and production phase, the system was described by the AMBER14SB forcefield[18] supplemented with Parmbsc1[19] and PET parameters using periodic boundary conditions and bond lengths correction by the LINCS algorithm[20] allowing to integrate the equations of motion according to Newton's 2$^{nd}$ law of motion every 2 fs using the leap-frog stochastic dynamics integrator.[21] Computational demand was reduced by utilization of the particle mesh Ewald method[22,23] for calculation of electrostatic interactions and by limiting the short-range non-bonded interactions to 12 Å.

Data was collected at least every 20 ps and analyzed using tools implemented in GROMACS[8] as well as visual molecular dynamics (VMD) software,[24] PyMOL[25] and Python 3.[26] Conformational clustering was performed using the Daura algorithm[27] with an RMSD cut-off of 2 Å. The EPS for the enzymes and PET was obtained using the Poisson-Boltzmann equation[28] as implemented in the adaptive Poisson-Boltzmann solver (APBS).[29]

#### 1.3 Enzyme-only simulations

The initial structures of LCC (PDB: 4EB0[30]) and PES-H1 (PDB: 7CUV,[10] chain B) were prepared by removing water, additives and additional enzyme chains in dimeric structures. Mutations were introduced using PyMOL to obtain the starting structures for the $LCC^{IG}$ variant (F243I and Y127G), the $LCC^{ICCG}$ variant ($LCC^{IG}$ including D238C and S283C mutations) and the PES-$H1^{FY}$ variant (L92F and Q94Y). The charge state at pH 7 of titratable amino acids was determined using Propka 3.4[31,32] yielding a net charge of +5 for LCC and $LCC^{IG}$, +6 for $LCC^{ICCG}$ and −5 for PES-H1 and PES-$H1^{FY}$, respectively. After energy minimization and equilibration, the enzymes were shortly simulated at 300 K (26.85°C) to check their stability and further relax the structures. Conformational clustering with a cutoff of 1 Å was applied using the Daura algorithm[27] to obtain the most populated cluster structure as a starting structure for subsequent simulations.

#### 1.4 Preparation of PET bulk

The initial *bulk.pdb* structure taken from Cruz-Chu *et al.*[33] consists of 245 PET chains, each including nine units with neutral carboxy termini. To decrease computational demand, the bulk was reduced to 100 PET chains and solvated, and this structure relaxed by MD simulations in three steps, which reduced the box size from 1680 $nm^3$ to 1040 $nm^3$. For each step, an MD simulation was conducted for 100 ns in absence of ions. A cubic box matching the size of the PET bulk was used together with semiisotropic pressure coupling to mimic an infinite PET surface when applying periodic boundary conditions. Conformational clustering of the PET bulk with a 3 Å cutoff yielded the most populated cluster as the starting structure for the respective following simulation, which finally included an enzyme.

#### 1.5 Simulating enzymes with PET bulk

The LCC and PES-H1 were subjected to MD simulations in a system with PET bulk to sample the initial adsorption to PET. The same x and y cubic box boundaries as for the PET bulk simulations as well as semiisotropic coupling in x and y direction were chosen to mimic an infinite PET surface under application of periodic boundary conditions. For each simulation, the box size in z direction was increased to make space for the incorporation for the enzyme, which was placed at least 10 Å minimum distance above the PET bulk in a random orientation using PyMOL.[25] Six different starting conformations were generated per enzyme to reduce the risk of a biased initial contact site and to increase statistical significance. An additional energy minimization was performed *in vacuo* to relax the structure prior to solvation. Simulations were carried out for 1.5 µs and the three most promising of the six simulations per enzyme, characterized by a minimum distance below 12 Å between the catalytic serine and PET, were elongated to 2 µs (LCC: $1^{st}$, $2^{nd}$, $4^{th}$; PES-H1: $1^{st}$, $2^{nd}$, $3^{rd}$ MD run), yielding a total simulation time of 10.5 µs per enzyme.

#### 1.6 Triple-mutant MD simulations

Variants of $LCC^{IG}$ and PES-$H1^{FY}$ were generated using PyMOL[25] to analyze the influence of charge and polarity on PET binding: T11V/S13A/S14A, S13A/R19E/R75E and Q26L/T27V/T28V for PES-$H1^{FY}$, and R143E/S145A/R151E, L66A/S67A/S69A and T211V/S216A/N239L for $LCC^{IG}$. As a starting conformation, we extracted the enzyme–PET bulk configuration in which PET was closest to these mutation sites from the preceding 1.5 µs MD simulations of the WT enzymes. In addition to the $LCC^{IG}$ and PES-$H1^{FY}$ triple mutants, we also simulated their respective parent enzyme, i.e., $LCC^{IG}$ and PES-$H1^{FY}$. All of these simulations were performed in triplicate to compare their interaction with PET and

PET contact progression with statistical significance. The box size was set according to the x and y boundaries of the extracted snapshots to fit the PET bulk conformation, but reduced in z direction to decrease computational demand (which is possible since the enzymes are already adsorbed on the PET surface here). An additional energy minimization was performed *in vacuo* prior to solvation and addition of ions. The pressure was scaled semiisotropically to allow for an infinite PET bulk surface under application of periodic boundary conditions. Simulations were performed for 100 ns each.

#### 1.7 Hamiltonian replica exchange MD (HREMD) simulations

In the MD simulations, adsorption of LCC and PES-H1 onto the PET bulk was observed with high retention near the catalytic site, but without PET penetrating into the binding cleft. To sample the entrance to the binding cleft, conformations with at most 5 Å distance between the catalytic serine and PET were extracted from the MD simulations that had been elongated to 2 µs, in which long retention of PET close to the active site took place. This yielded one PET pose for PES-H1 (1st run, 3.25 Å after 25.6 ns) and two poses for LCC (1st run, 1.59 Å after 1,264.4 ns; 4th run, 1.69 Å after 58.8 ns), which were used for further study. However, only the enzyme and the one PET chain closest to the catalytic serine were kept, which was further reduced from nine to the five PET units closest to the catalytic serine to reduce the computational demand of the following HREMD simulations. To increase the number of starting configurations to three per enzyme, the two LCC-PET poses were also used for PES-H1 by replacing LCC with PES-H1 after superposition of the enzymes, and vice versa the original PES-H1-PET configuration was copied to LCC, denoted as 1stP (from the $1^{st}$ PET bulk simulation with PES-H1) and 1stL and 4thL (from the $1^{st}$ and $4^{th}$ bulk simulations with LCC). The simulations were prepared for the WT enzymes and variants listed in Tab. S1 using the standard equilibration MD protocol described above. This was followed by an additional simulation step, which was conducted for 10 ns using the same conditions as for NpT equilibration but without position-restraining the enzyme to relax the protein structure and ensure system stability prior to the HREMD production run, which was conducted for 115 ns per replica using GROMACS version 2016.4 patched with the PLUMED 2.4.6 plugin.[8,34,35] Here, the v-rescale thermostat[13] is used instead of the Nosé-Hoover thermostat[15,16] as it is recommended for the HREMD method.[35] Six replicas per HREMD simulation were used, where the Hamiltonian of PET's non-bonded interactions was perturbed by applying $\lambda$ scaling factors of 1.00, 0.87, 0.76, 0.66, 0.57, and 0.50, corresponding to temperatures ranging from 300 to 600 K. Exchange rates of approx. 30% were obtained with exchanges between replicas being attempted every 0.2 ps. For some of the variants, multiple HREMD simulations were performed, which were initiated with different initial velocities. In total, we run 37 and 31 HREMD simulations corresponding to 25.53 and 21.39 µs of sampling for variants of LCC and PES-H1, respectively (Tab. S1).

Besides the HREMD production run, conventional MD production runs were performed at 303 K and 343 K for the three starting structures 1stP, 1stL, and 4thL with only one PET chain (of five units), as well as one simulation for each starting structure including the whole PET bulk at 343 K. These extra MD simulations were done to ensure that the HREMD results obtained are not a consequence of reducing the PET to one chain or the HREMD settings and that they are reproducible at higher temperatures and with the inclusion of a PET bulk. The same simulation conditions as in the HREMD simulations were applied for the systems including one PET chain, while the conditions from the initial PET bulk simulations were used for the systems including PET bulk.

Table S1: Variants studied by HREMD simulations with the starting structures 1stP (taken from 1st PET-bulk simulation of PES-H1), 1stL, and 4thL (taken from the 1st and 4th PET-bulk simulations of LCC). The number of HREMD simulations performed per variant and starting structure as well as the total number of simulations are given. For both enzymes, 16 mutation positions were analyzed.

| **LCC** | **1stP** | **1stL** | **4thL** | **total** | **PES-H1** | **1stP** | **1stL** | **4thL** | **total** |
|---|---|---|---|---|---|---|---|---|---|
| WT | 1 | 1 | 3 | 5 | WT | 2 | 1 | 1 | 4 |
| $LCC^{IG}$ | 2 | 2 | 1 | 5 | PES-$H1^{FY}$ | 1 | 1 | 1 | 3 |
| T96A | 0 | 0 | 1 | 1 | T63G | 0 | 0 | 1 | 1 |
| S101A | 0 | 0 | 1 | 1 | S68A | 0 | 0 | 1 | 1 |
| | | | | | S68D | 0 | 0 | 1 | 1 |
| | | | | | S68F | 0 | 0 | 1 | 1 |
| | | | | | S68I | 0 | 0 | 1 | 1 |
| | | | | | S68K | 0 | 0 | 1 | 1 |
| | | | | | S68Q | 0 | 0 | 1 | 1 |
| T121A | 0 | 0 | 1 | 1 | D93A | 0 | 0 | 1 | 1 |
| R124A | 0 | 1 | 0 | 1 | R91A | 0 | 0 | 1 | 1 |
| F125L | 0 | 0 | 1 | 1 | R98A | 0 | 0 | 1 | 1 |
| D126A | 0 | 0 | 2 | 2 | H129A | 0 | 0 | 1 | 1 |
| R131A | 0 | 0 | 1 | 1 | T157G | 0 | 1 | 0 | 1 |
| H164A | 0 | 0 | 1 | 1 | L175A | 0 | 0 | 2 | 2 |
| T192G | 1 | 0 | 0 | 1 | T177G | 2 | 0 | 1 | 3 |
| T211G | 0 | 0 | 2 | 2 | H184S | 1 | 0 | 1 | 2 |
| Q217A | 0 | 0 | 1 | 1 | S207A | 0 | 0 | 1 | 1 |
| H218A | 0 | 0 | 1 | 1 | L209A | 1 | 0 | 0 | 1 |
| S241G | 0 | 1 | 1 | 2 | N212A | 0 | 1 | 1 | 2 |
| N246A | 3 | 0 | 0 | 3 | T213A | 0 | 0 | 1 | 1 |
| **total** | | | | 32 | **total** | | | | 31 |

#### 1.8 Free energy surface (FES)

Two-dimensional FESs were generated for a temperature of 303 K using the HREMD data. The two collective variables along which the free energies $\Delta G$ were projected were calculated using tools implemented in GROMACS: *gmx select* and *gmx mindist* for the SER–C distance and *gmx energy* as well as *gmx rerun* to obtain the total interaction energy $E_{\text{int}}$ between PET and specific residues consisting of polar Coulomb (Coul) and hydrophobic Lennard-Jones (LJ) contributions. The multistate Benett acceptance ratio (MBAR) method as implemented in the Python 3 package *pymbar*[36–38] was used to construct the FES histogram. To reduce noise and for representing the FESs as smooth contour plots, a Gaussian filter was used as implemented in the Python module *scipy.ndimage*.

#### 1.9 Principal component analysis (PCA)

PCA was performed using the minimum distance between the post-entry productively bound PET ester carbon and all $C_\alpha$ atoms that were located at least once within 10 Å of that PET carbon during entry of the PET and the first 1 ns of its inside phase. The resulting data from both enzymes were merged, taking into account corresponding $C_\alpha$ atoms that were identified by superimposing both enzyme structures, resulting in a total of 49 $C_\alpha$ positions that were used in the PCA. The dataset of distances between these $C_\alpha$ atoms and PET, as described above, was subjected to PCA by using the Python 3 module *PCA* as implemented in the *decomposition* package of the *scikit learn* library (*sklearn.decomposition.PCA*).[39] The resulting principal components (PCs) are thus identical for both enzymes, yet the individual datasets per enzyme were used for constructing FESs at 303 K along the two main PCs using an inhouse Python 3 script, which calculates $\Delta G$ based on the probability distribution $P$ along the order parameter $R$, here the two PCs (Eq. 1).

$$\Delta G(R) = -k_B T \left[\ln P(R) - \ln P_{\max}(R)\right] \tag{1}$$

The resulting FESs were further analyzed by identifying productive and unproductive states in the PC space and by projecting individual entry events onto the FESs.

#### 1.10 Enzyme production and purification

PES-H1$^{FY}$ and LCC$^{ICCG}$ variants were produced using pET28a(+) and pET26b(+) as vectors, respectively. Genes encoding PES-H1$^{FY}$ and LCC$^{ICCG}$ have been synthesized and codon optimized as previously described.[10] Chemically competent *E.coli* BL21 DE3 cells were transformed with the plasmids and overnight cultures containing 50 µg/ml kanamycin were inoculated from glycerol stocks. For the expression, 250 ml of ZYM-5052 autoinduction medium containing 50 µg/ml kanamycin were added into a 1 L baffled flask and 2.5 ml of overnight culture were used for the inoculation. Cell cultures were incubated at 37 °C; 120 rpm for approximately 2 h and subsequently the temperature was reduced to 21 °C. After 21 h or 23 h of incubation for PES-H1$^{FY}$ and LCC variants, respectively, cells were centrifuged at 12,000 g for 15 min, and pellets were resuspended in 4 ml/g cell pellet of lysis buffer (50 mM $Na_2HPO_4$, 300 mM NaCl, pH 8.0) and lysed using ultrasonication (2x 3 min at 50% power, 50% amplitude). After subsequent centrifugation at 12,000 g for 30 min, the enzymes were purified using cobalt-ion affinity chromatography (ROTI®Garose-His/Co Beads, Carl Roth, Karlsruhe, Germany). The supernatant was applied onto a gravity flow column and washed with either 20 mM or 50 mM imidazole (pH 8.0). The proteins were eluted with an elution buffer containing 50 mM $Na_2HPO_4$, 300 mM NaCl and 250 mM imidazole (pH 8.0). The imidazole was removed with a PD-10 desalting column (Cytiva, Marlborough, Massachusetts, USA) changing the buffer to 50 mM $Na_2HPO_4$ (pH 8.0) and proteins were concentrated with 10 kDa molecular weight cutoff centrifugal concentrators (Satorius, Goettingen, Germany). The protein concentration was determined with a Nanodrop1000 spectrophotometer.

#### 1.11 Site directed mutagenesis

Single Mutations were introduced using the Q5® Site Directed Mutagenesis Kit (New England Biolabs GmbH, Ipswich, Massachusetts, USA). Oligonucleotides used in the polymerase chain reaction (PCR) are listed in Table S2. The successful incorporation of the mutation was confirmed by Sanger sequencing (Microsynth AG, Balgach, Switzerland).

Table S2: Oligonucleotides used for site directed mutagenesis

| **Mutation** | **Forward Primer** | **Reverse Primer** |
|---|---|---|
| LCC$^{ICCG}$ H164A | GGTGGCGGGT**gcg**AGCATGGGCGG | GCCAGACGGTTGGCATCC |
| LCC$^{ICCG}$ T192G | GCCGTGGCAT**ggc**GATAAAACCTTT AA-CACCAGCGTGC | GTCAGCGGCACCGCCGCT |
| LCC$^{ICCG}$ T211G | GGAAGCCGAT**ggc**GTGGCGCCGG | GCGCCAACAATCAGCACC |
| LCC$^{ICCG}$ S241G | GTGCAACGCC**ggc**CATATTGCGC | AGTTCAACATAAACTTTC GGCGTGGTG |
| LCC$^{ICCG}$ N246A | TATTGCGCCG**gcg**AGCAATAACGCGG | TGGCTGGCGTTGCACAGT |
| PES-H1$^{FY}$ S68A | CGGTCAGGAA**gcg**ATTGCCTGGCTG GGCCC | GCGGTAAAGCCCGGGCTG |
| PES-H1$^{FY}$ R91A | TACCATCACC**gcg**TTTGATTATCCG GAC | TCAATGGTAATCACCACAAAG |
| PES-H1$^{FY}$ D93A | CACCCGTTTT**gcg**TATCCGGACAG | ATGGTATCAATGGTAATCAC |
| PES-H1$^{FY}$ L175A | TGGCGCGCAG**gcg**GATACCATTGCGC | ACCACCAGCGTCGGCGTG |
| PES-H1$^{FY}$ N212A | TCTGGTCAGC**gcg**ACGCCGGATACC AC-CACC | TGGCTGGCACCGCGCAGT |

#### 1.12 Enzymatic hydrolysis of PET film

The PET-film (Goodfellow GmbH, product number ES301445) was cut into 1x 2 cm pieces (∼60 mg), which were washed once with ethanol, afterwards with $dH_2O$ and were dried at room temperature. Degradation of PET was performed with 1x 60 mg PET film in a 2 ml reaction vessel with an enzyme load of either 1 mg enzyme/g PET or 0.5 mg enzyme/g PET and 1 M potassium phosphate-buffer was added to a total volume of 1.5 ml. The reaction mixtures were incubated at 70 °C on a ThermoMixer C (Eppendorf, Hamburg, Germany) for 24 h at 1000 rpm. Each reaction was performed at least in duplicates. After 24 h, the remaining PET-films were washed in a 0.1% SDS solution, $dH_2O$ and absolute ethanol and subsequently dried at 50 °C for 24 h. The weight loss through PET degradation was determined gravimetrically.

#### 1.13 Kinetic analysis

For determining the kinetic parameters, a turbidimetric assay was applied: A mixture of 3.75 ml ROTIPHORESE® gel 30, 3 ml potassium phosphate buffer (1 M, pH 8.0), 120 µl tetramethylethylenediamine (TEMED) and PET-NP suspension (1 mg·ml$^{-1}$) was prepared and filled up to a final volume of 15 ml with dd$H_2O$. PET-NP were prepared as previously described.[40–42] A total volume of 50 µl enzyme mix was prepared by combining 2 µl of a 40% (w/v) solution of ammonium persulfate (APS) with 48 µl of the enzyme. This mixture was then mixed with 150 µl of the solution containing the substrate and acrylamide. The turbidimetric assay was carried out in a microtiter plate at a temperature of 70 °C for a duration of 40 min for each variant at least in triplicate. The substrate load concentrations varied from 0-1.5 mg/ml while keeping a constant enzyme concentration of 0.7 µM for the conventional Michaelis–Menten ($^{\mathrm{conv}}$MM) setting (Eq. 2).

$$ {}^{\mathrm{conv}}v_0 = \frac{{}^{\mathrm{conv}}V_{\mathrm{max}}[S_0]}{{}^{\mathrm{conv}}K_{\mathrm{m}} + [S_0]} \tag{2} $$

The inverse Michaelis–Menten ($^{\mathrm{inv}}$MM) approach included enzyme concentrations varying from 0-6 µM while keeping a fixed PET-NP concentration of 0.375 mg/ml(Eq. 3).

$$ {}^{\mathrm{inv}}v_0 = \frac{{}^{\mathrm{inv}}V_{\mathrm{max}}[E_0]}{{}^{\mathrm{inv}}K_{\mathrm{m}} + [E_0]} \tag{3} $$

The turnover rate constant $k_{cat}$ and the catalytic efficiency $^{\mathrm{mass}}\eta$ can be obtained from the $^{\mathrm{conv}}$MM (Eq. 4 and 5).

$$ k_{cat} = \frac{{}^{\mathrm{conv}}V_{\mathrm{max}}}{[E_0]} \tag{4} $$

$$ {}^{\mathrm{mass}}\eta = \frac{k_{cat}}{{}^{\mathrm{conv}}K_{\mathrm{m}}} \tag{5} $$

For a better comparability when targeting large, insoluble substrates, the catalytic efficiency is also assessed in terms of molarity (${}^{\text{molar}}\eta$) instead of mass (${}^{\text{mass}}\eta$). This is not possible considering only the kinetic parameters obtained from ${}^{\text{conv}}$MM since ${}^{\text{conv}}K_M$ considers only the used substrate load in terms of mass ${}^{\text{mass}}[S_0]$ and not in terms of molarity as in the ${}^{\text{inv}}$MM approach. The accessible attack sites on the PET surface (${}^{\text{molar}}[S_0]$) can be obtained by defining the density of attack sites per gram PET ($\Gamma_{\text{attack}}$) (Eq. 6).

$$ {}^{\text{molar}}[S_0] = {}^{\text{mass}}[S_0] \cdot \Gamma_{\text{attack}} \tag{6} $$

$\Gamma_{\text{attack}}$ can be obtained by taking the ${}^{\text{inv}}$MM parameters into account (Eq. 7), which then also allows to estimate a molar ${}^{\text{molar}}K_{\text{m}}$ for the ${}^{\text{conv}}$MM setting (Eq. 8) and, thus, the ${}^{\text{molar}}\eta$ (Eq. 9).[43,44]

$$ \Gamma_{\text{attack}} = \frac{{}^{\text{inv}}V_{\text{max}}/[S_0]}{{}^{\text{conv}}V_{\text{max}}/[E_0]} \tag{7} $$

$$ {}^{\text{molar}}K_{\text{m}} = {}^{\text{conv}}K_{\text{m}} \cdot \Gamma_{\text{attack}} \tag{8} $$

$$ {}^{\text{molar}}\eta = \frac{k_{cat}}{{}^{\text{molar}}K_{\text{m}}} = \frac{{}^{\text{mass}}\eta}{\Gamma_{\text{attack}}} \tag{9} $$

#### 2 Supplementary Figures

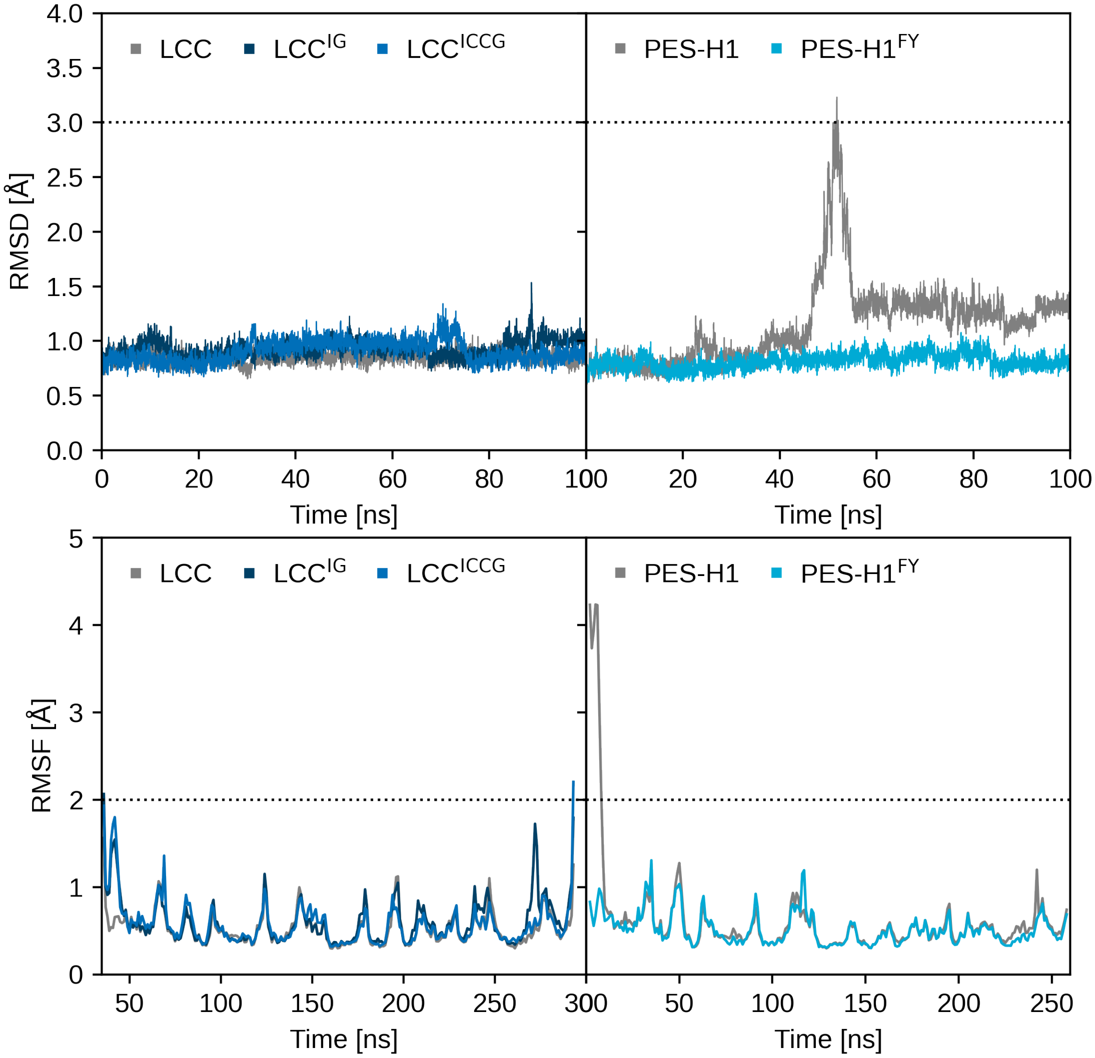


Figure S1: RMSD (top) and RMSF (bottom) of LCC, LCC$^{IG}$ and LCC$^{ICCG}$ (gray, darkblue and blue, left) and PES-H1 and PES-H1$^{FY}$ (gray and lightblue, right). We define an RMSD difference of 3 Åwith respect to the starting structure of the simulation as a significant structural difference, which is only observed once for PES-H1 at around 50 ns. This structural change results from structural flexibilities in the first ten disordered N-terminal residues, which can be recognized by the large RMSF values in this region. The RMSF specifies the flexibility per residue, whereby an RMSF below 2 Å defines a rigid residue and above this value a flexible one. Overall, the structures of all four proteins were consistently rigid throughout the simulations.

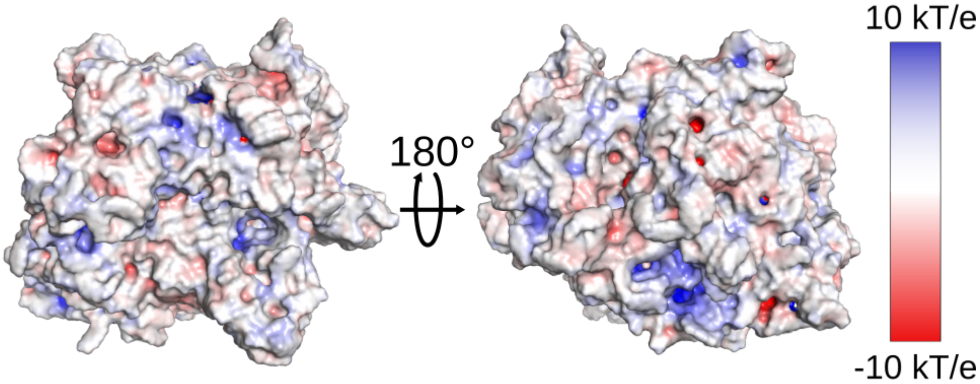


Figure S2: The EPS of the PET bulk used in this study is shown from two perspectives with a positive EPS colored in blue and a negative colored in red, while hydrophobic areas are shown in white.

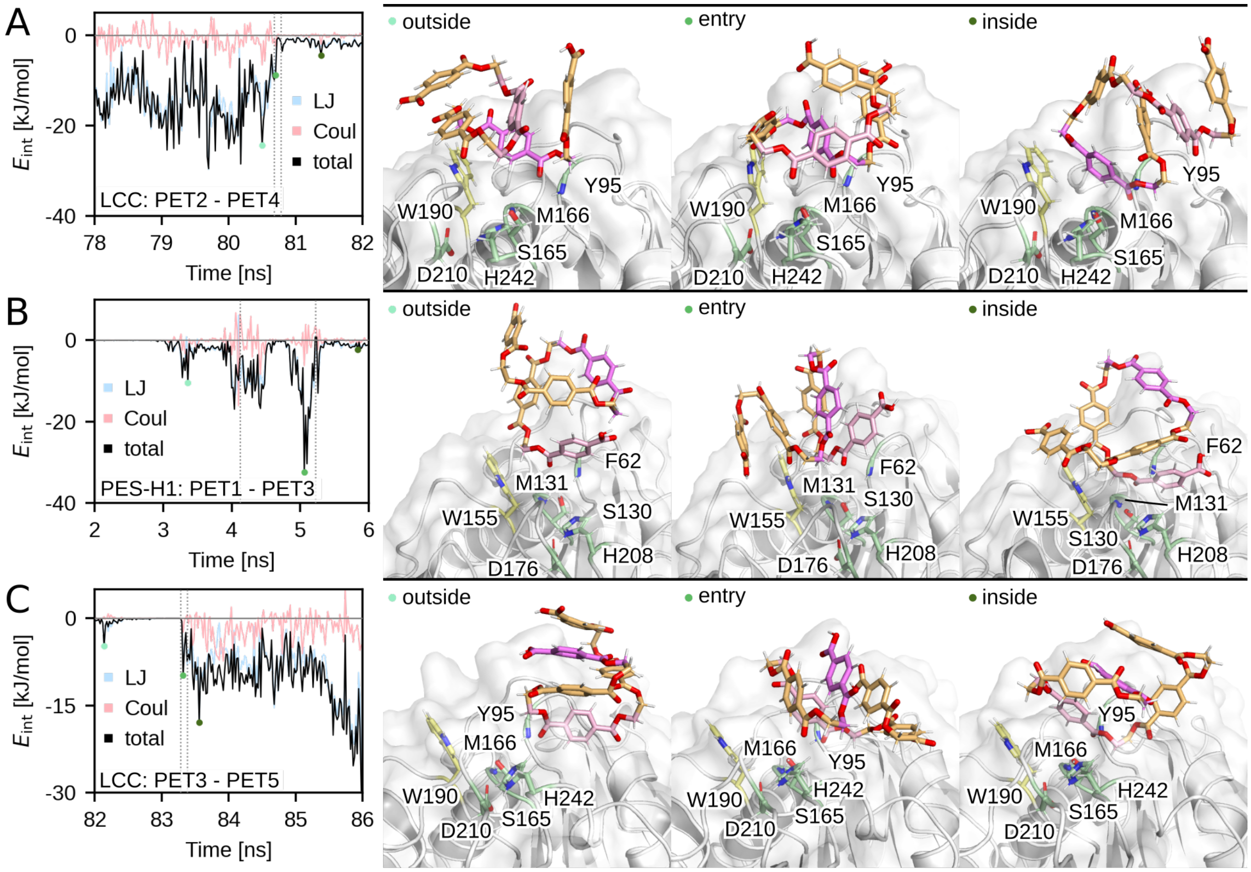


Figure S3: Examples of intramolecular PET interactions before PET entry into the binding site (A, LCC, PET2–PET4 interactions), during entry (B, PES-H1, PET1–PET3), and after entry (C, LCC, PET3–PET5). The left plots show the evolution of these interaction energies, $E_{\mathrm{int}}$, which comprise polar Coulomb (Coul) and hydrophobic Lennard-Jones (LJ) interactions, during 4 ns snippets of the entry process. The actual entry phase is indicated by vertical dotted lines. Structures for each of the three entry events are shown on the right-hand side, with a snapshot from each phase (outside, entry, inside) being presented. The times at which these snapshots were sampled are indicated by green-colored spheres in the plots on the left. In the structures, the catalytic triad and the oxyanion hole are shown in green, the 'wobbling' tryptophan in yellow, PET in orange, and the two interacting PET units in light pink (lower residue ID) and dark pink (higher residue ID).

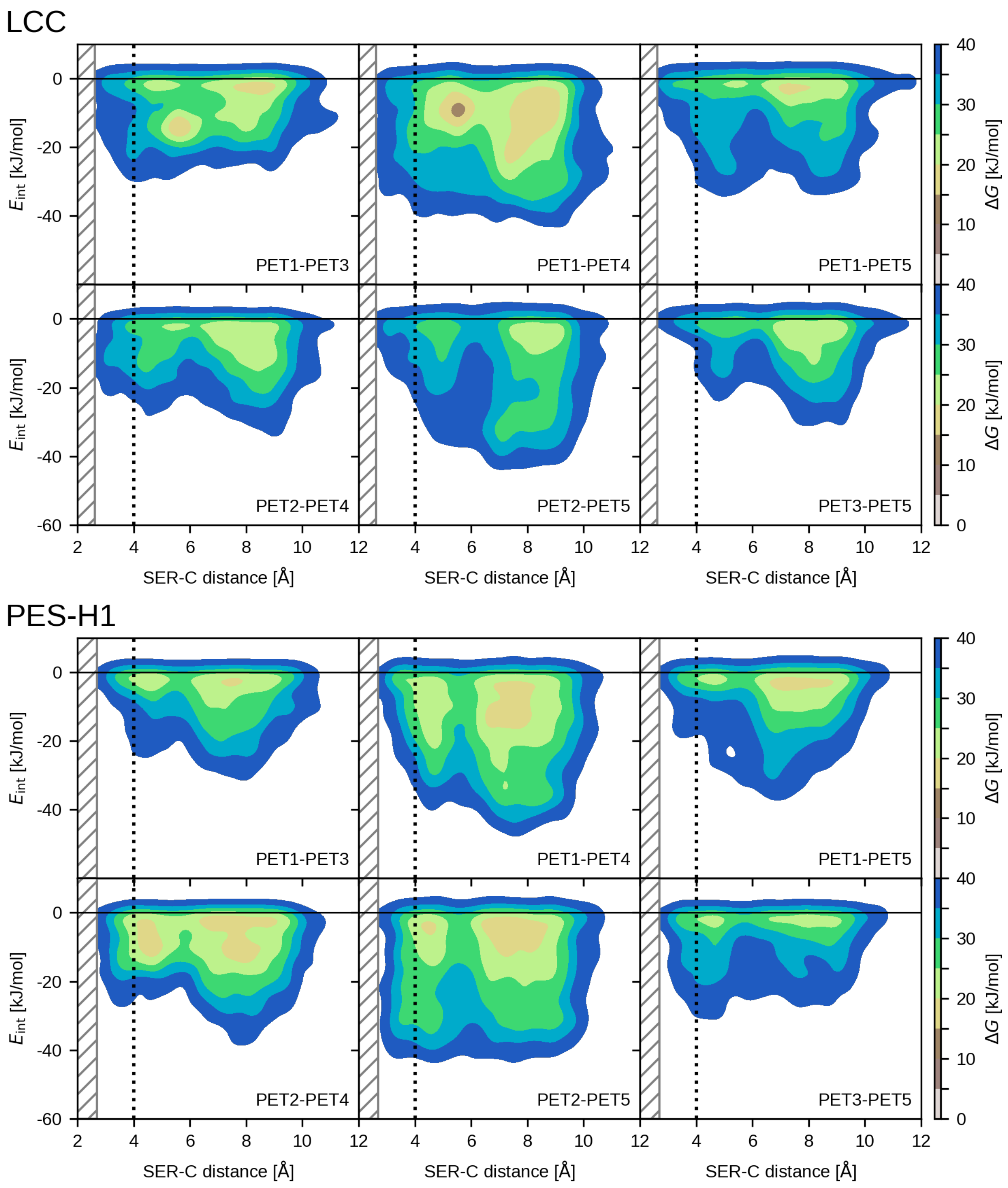


Figure S4: Free energy surfaces of PET–PET interactions over the SER–C distance sampled during LCC (top) and PES-H1 (bottom) simulations. The area with a minimum SER–C distance <4 Å harbors productive poses. The gray hatched area indicates distances that were not sampled, resulting in a truncated FES.

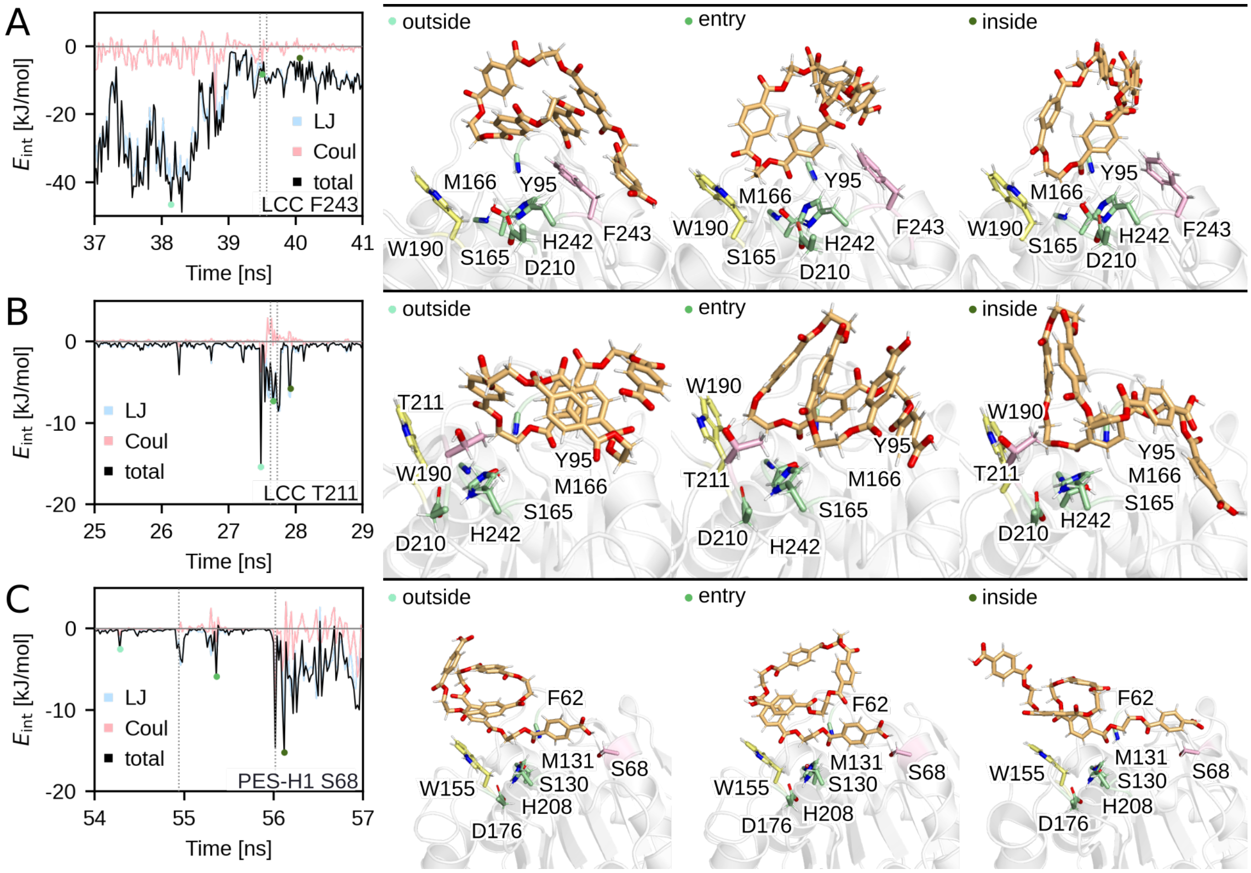


Figure S5: Examples of residues interacting with PET before PET entry into the binding site (A, with LCC F243), during entry (B, with LCC T211), and after entry (C, with PES-H1 S68). The left plots show the evolution of these interaction energies, $E_{\text{int}}$, which comprise polar Coulomb (Coul) and hydrophobic Lennard-Jones (LJ) interactions, during 4 ns snippets of the entry process. The actual entry phase is indicated by vertical dotted lines. Structures for each of the three entry events are shown on the right-hand side, with a snapshot from each phase (outside, entry, inside) being presented. The times at which these snapshots were sampled are indicated by green-colored spheres in the plots on the left. The catalytic triad and oxyanion hole are shown in green, the 'wobbling' tryptophan in yellow, PET in orange, and the interacting residue in pink.

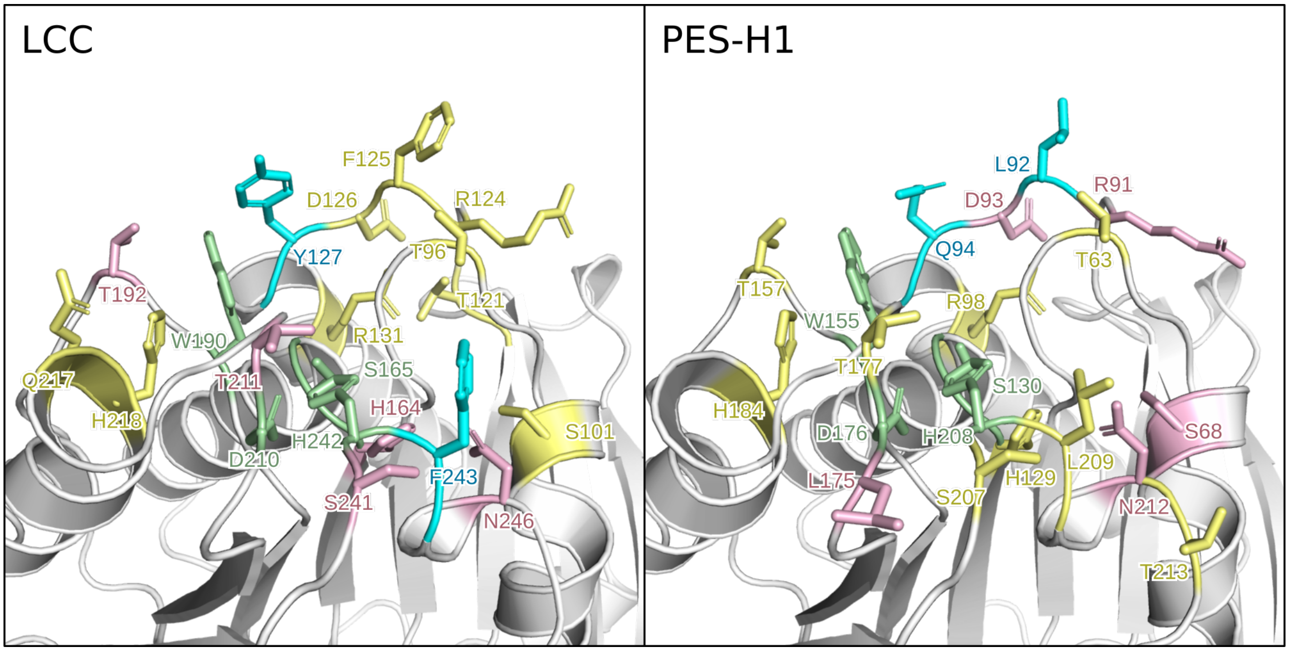


Figure S6: Mutation sites of variants addressed using the FES approach. The catalytic triad residues as well as the 'wobbling' tryptophan are shown in green, the LCC$^{IG}$ and PES-H1$^{FY}$ mutation sites in cyan, the experimentally evaluated mutation sites in red, and all other mutation sites in yellow.

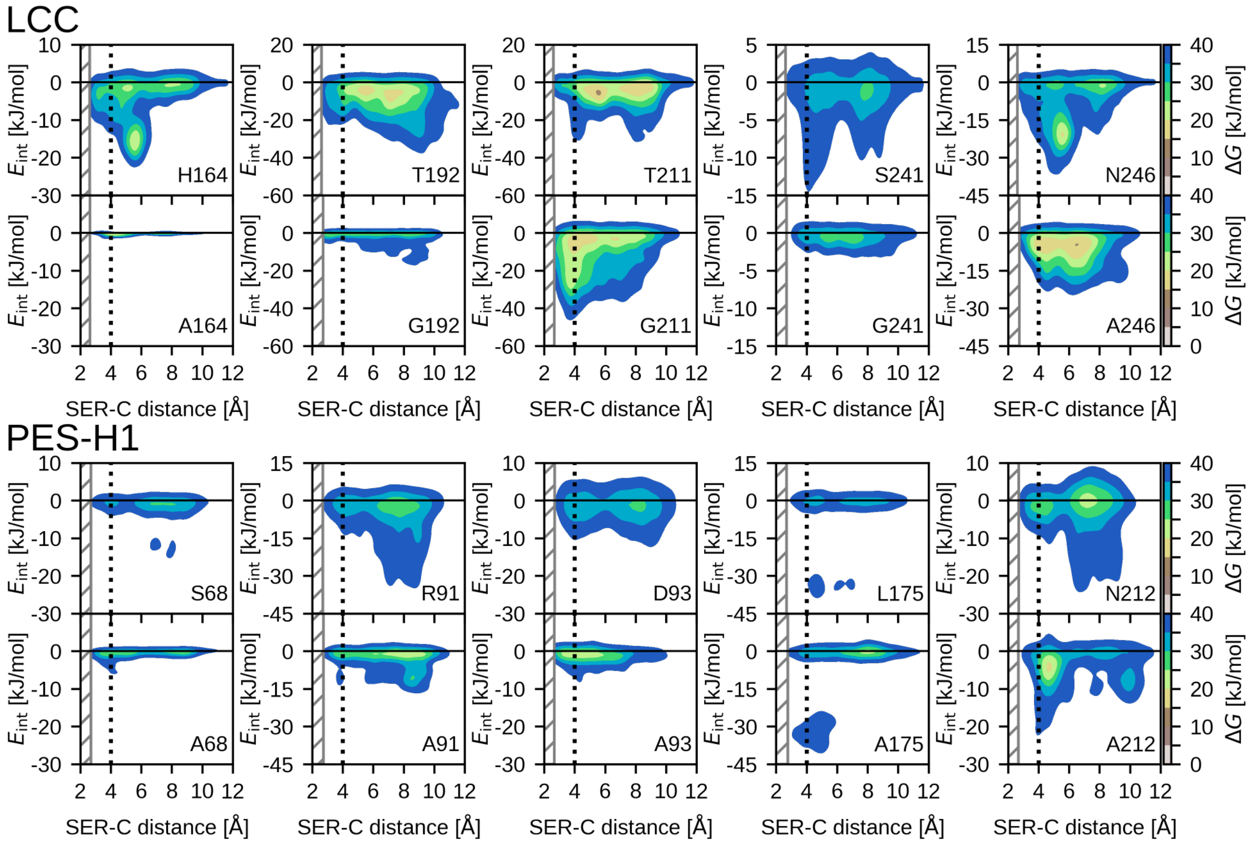


Figure S7: Free energy surfaces of PET–residue interactions over the SER–C distance sampled during LCC and PES-H1 and corresponding variant simulations. The top rows present the FES for WT residues and the bottom row for mutated residues. The area with a minimum SER–C distance <4 Å harbors productive poses. The gray hatched area indicates distances that were not sampled, resulting in a truncated FES.

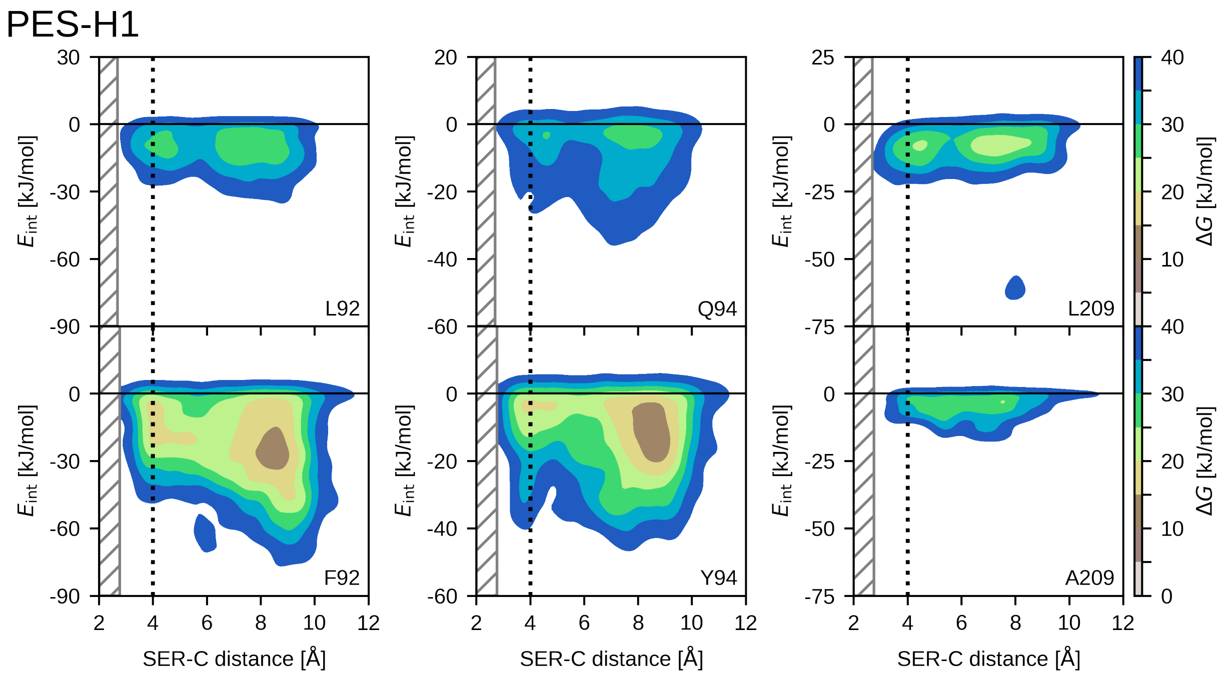


Figure S8: Free energy surfaces of PET–residue interactions at position 68 over the SER–C distance sampled during PES-H1 and corresponding PES-H1$^{FY}$ variant simulations. The left FES shows the interaction of the WT S68 and the remaining ones of variants. The area with a minimum SER–C distance <4 Å harbors productive poses. The gray hatched area indicates distances that were not sampled, resulting in a truncated FES.

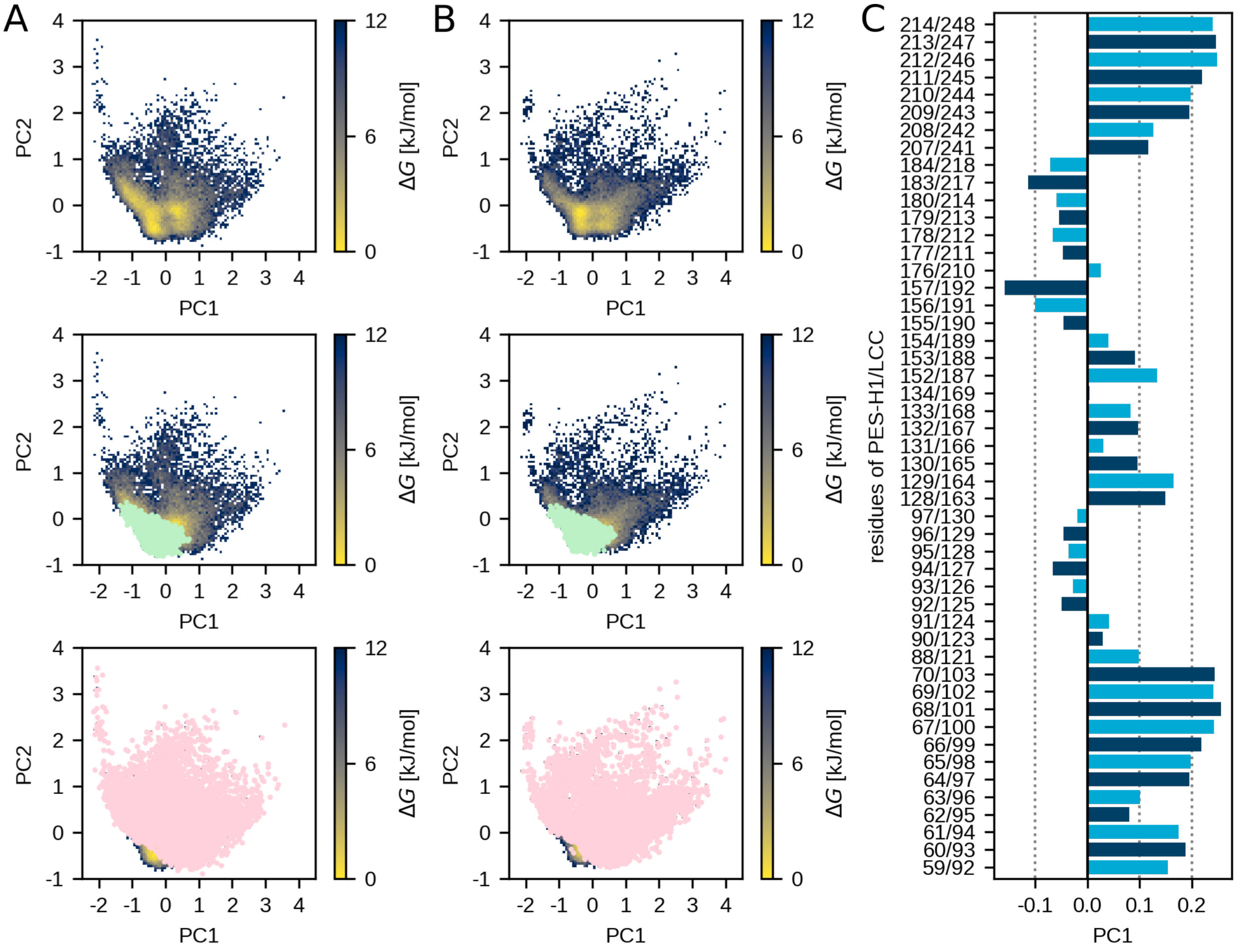


Figure S9: The FES over PC1 and PC2 of LCC (A) and PES-H1 (B), accompanied by the contribution profile of amino acid positions to PC1 (C). In the central figure of A and B, productive states of PET in the active site of each enzyme are highlighted in green, while the lower figure depicts unproductive states in red.

### 3 Supplementary Tables

Table S3: Maximum reaction rate $V_{\max}$ for both settings, ${}^{conv}$MM, ${}^{inv}$MM derived from the nonlinear fit to experimental data as well as ${}^{molar}\eta$ and $\Gamma_{attack}$.

| | ${}^{conv}\mathbf{V}_{\max}[\mathbf{min^{-1}}]$ | ${}^{inv}\mathbf{V}_{\max}[\mathbf{min^{-1}}]$ | ${}^{mass}\eta[\frac{\mathbf{L^2}}{\mathbf{g \cdot min \mu mol}}$ | $\mathbf{\Gamma}_{attack}\left[\frac{\mathbf{\mu mol}}{\mathbf{g}}\right]$ |
|---|---|---|---|---|
| LCC$^{ICCG}$ | 0.033 ±0.003 | 0.029 ±0.001 | 0.219 | 1.659 |
| LCC$^{ICCG}$ H164A | 0.022 ±0.003 | 0.035 ±0.002 | 0.335 | 2.931 |
| LCC$^{ICCG}$ T211G | 0.029 ±0.005 | 0.039 ±0.003 | 0.462 | 2.450 |
| LCC$^{ICCG}$ S241G | 0.038 ±0.003 | 0.029 ±0.002 | 0.217 | 1.433 |
| PES-H1$^{FY}$ | 0.062 ±0.006 | 0.088 ±0.003 | 0.226 | 2.646 |
| PES-H1$^{FY}$ S68A | 0.059 ±0.007 | 0.064 ±0.003 | 0.234 | 2.038 |
| PES-H1$^{FY}$ D93A | 0.028 ±0.004 | 0.072 ±0.007 | 0.079 | 4.860 |
| PES-H1$^{FY}$ N212A | 0.047 ±0.007 | 0.082 ±0.006 | 0.140 | 3.250 |
